# Distribution, assembly and mechanism of GluN1/GluN3A excitatory glycine receptors

**DOI:** 10.64898/2026.03.29.715060

**Authors:** Lizhen Xu, Marco De Battista, Kejie Yao, Jochen Schwenk, Lea Nehme, Lara Pizzamiglio, Anto Cerasale, Bernd Fakler, David Stroebel, Shujia Zhu, Pierre Paoletti

## Abstract

NMDA receptors play key roles in brain development, plasticity and diseases. While glutamate and glycine co-gated GluN2-containing NMDARs have been extensively characterized, little is known regarding GluN3A-containing NMDARs that form receptors gated by glycine only. Here, combining native purification, mass spectrometry, cryo-EM and electrophysiology, we provide key insights on the molecular logic of GluN3A-NMDARs. We demonstrate that native GluN3A receptors account for a sizeable fraction of total NMDARs, are enriched at extrasynaptic compartments in the adult brain, and assemble specifically as diheteromeric GluN1/GluN3A excitatory glycine receptors (eGlyRs) rather than as triheteromeric GluN1/GluN2/GluN3A receptors. The architecture of eGlyRs reveal splayed and loosely packed extracellular domains, strikingly different from ‘conventional’ GluN1/GluN2 receptors. Through back-and-forth structural and functional validations, we demonstrate how the combined effects of a weak ligand-binding domain (LBD) dimer interface and high intrinsic mobility of the N-terminal domains (NTDs) shape the atypical gating of eGlyRs. These findings illuminate GluN3A-NMDAR physiology and mechanism, with implications for neuronal signaling and pharmacology.

## Introduction

Excitatory neurotransmission and cellular signaling mediated by *N*-methyl-D-aspartate receptors (NMDARs) play critical roles in brain development, function and diseases^1,2^. Canonical NMDARs are heterotetrameric ligand-gated ion channels formed by two GluN1 subunits and two GluN2(A-D) subunits whose activation requires the simultaneous binding of glycine (on GluN1 subunits) and glutamate (on GluN2 subunits)^1–3^.

GluN1/GluN2 NMDARs typically display high Ca^2+^ permeability, strong voltage-dependent Mg^2+^ block, and are essential drivers of long-term synaptic plasticity underlying learning and memory^1,4^. The structure, gating mechanism and pharmacology of GluN1/GluN2 NMDARs have been extensively studied and characterized, providing a wealth of information on the molecular basis of glutamatergic excitatory neurotransmission^5–14^.

In contrast, much less is known regarding NMDARs composed of GluN3 subunits, of which there are two subtypes (GluN3A and GluN3B)^15,16^ that, like GluN1, also bind glycine^3,17–19^. GluN3A expression peaks in the juvenile brain and is maintained at sustained levels in the adult brain, suggesting a life-long importance of GluN3A-receptors in neural function^20,21^. In contrast, GluN3B expression is much more limited throughout development, with little or no expression in the forebrain^21,22^. Genetic evidence implicates GluN3A subunits in brain maturation and synapse development^20,21,23,24^. Moreover, alterations of GluN3A expression in humans are linked to diverse neuropsychiatric conditions, such as schizophrenia, bipolar disorders and epilepsy^20,21^, making GluN3A-receptors an attractive target for therapeutic intervention. Traditionally, roles of GluN3A were interpreted as having direct influence on conventional GluN2-NMDARs via the formation of GluN1/GluN2/GluN3A triheteromeric NMDARs. In these receptors, the GluN3A subunit serves as a dominant negative subunit, dampening channel gating by glutamate and glycine and calcium permeation^20^.

However, this paradigm was recently challenged by the realization that GluN3A subunits can readily assemble with GluN1 in diverse neuronal populations to form functional GluN1/GluN3A receptors unresponsive to glutamate and solely gated by glycine^25–32^. These excitatory glycine receptors (eGlyRs), first described in recombinant systems^25,33–35^, have now been found in several brain regions, where they act as sensors of extracellular glycine to control neuronal excitability^21,27^. The eGlyRs are strongly enriched and behaviorally active in key brain regions encoding emotional states and stress responses, such as medial habenula^26^, basolateral amygdala^27^ and ventral hippocampus^31^. In contrast to conventional GluN2-NMDARs, eGlyRs display low calcium permeability and minimal blockade by extracellular magnesium ions^21,33^. These receptors thus represent an emerging glycine-mediated signaling system in the CNS with high biomedical potential.

The discovery of eGlyRs obliges re-interrogation of the relevance of triheteromeric GluN1/GluN2/GluN3A, whose very existence is currently disputed^20,21,31,36^. Moreover, because GluN1/GluN3A NMDARs (i.e. eGlyRs) differ fundamentally from conventional GluN2-NMDARs in both their expression pattern and their operational modalities, there is considerable interest in better understanding how eGlyRs compare to GluN2-NMDARs. In this work, combining biochemical and functional approaches on native receptors as well as structural and mutagenesis analysis on recombinant receptors, we provide key insights into the molecular biology of GluN3A-receptors. We show that GluN3A subunits assemble specifically as GluN1/GluN3A diheteromers and are abundant in the juvenile brain. In the adult brain, eGlyRs are maintained at a significant level, representing a sizeable fraction of all NMDARs with a strong enrichment in non-synaptosome compartments. We also determine the structure of full-length GluN1/GluN3A receptors captured in various conformational and ligand-bound states. Through back-and-forth interplay between cryo-EM structural data and electrophysiological validation, we further identify key structural regions, including the LBD dimer interface and NTD-LBD linkers, that shape the unique gating and ion permeation properties of GluN1/GluN3A receptors. Through targeted residue substitution in these regions, we are able to directly manipulate the gating and block the fast desensitization characteristic of these glycine-gated receptors. Overall, our results provide a long-awaited quantitative and structural framework for GluN3A-NMDAR molecular physiology, highlighting key differences with conventional GluN2-NMDARs and other ionotropic glutamate receptor (iGluR) family members. These findings have direct implications for our understanding of how endogenous and pharmacological agents modulate eGlyRs and neuronal function in physiological and pathological conditions.

## Results

### Developmental pattern and subcellular localization of native GluN3A receptors

We first aimed to characterize the developmental distribution and subcellular expression patterns of GluN3A and other NMDAR subunits. For that purpose, we performed liquid chromatography-mass spectrometry (LC-MS)-based proteomic quantification on native whole brain extracts from postnatal day 10 (P10) and 8-week-old mice. We focused on three NMDAR-enriched samples termed as postsynaptic density (PSD), non-PSD and non-synaptosome. The first fraction refers to the PSD from the postsynaptic membrane, and the two other fractions enriched using the GluN1 antibody Mab_4F11_^10^ to detergent-solubilized non-PSD synaptic membranes and non-synaptosome membranes, respectively (see Methods for details; Fig. 1a). Mass spectrometry analysis showed that GluN1 was consistently detected at ∼50% across all three fractions (Fig. 1a), confirming its obligatory role in NMDAR assembly^2^. The GluN2B was the predominant GluN2 subunit in P10 brain and decreased in abundance in adulthood, accompanied by a reciprocal increase in GluN2A (Fig. 1a), aligning with the well characterized GluN2B to GluN2A developmental ‘switch’^2,37–39^. In the adult brain, GluN2A and GluN2B were present in comparable and substantial proportions (Fig. 1a), consistent with the subunit composition of native NMDARs^10,40^. Notably, GluN2B was abundant in all fractions (with the highest levels in the PSD fraction), supporting a non-segregation rule of GluN2B between synaptic and extrasynaptic compartments^39,41^. The GluN2D subunit was also detected at and outside synapses, but in lower in abundance, in line with its expression mainly restricted to GABAergic interneurons^1,42^.

**Figure 1.**
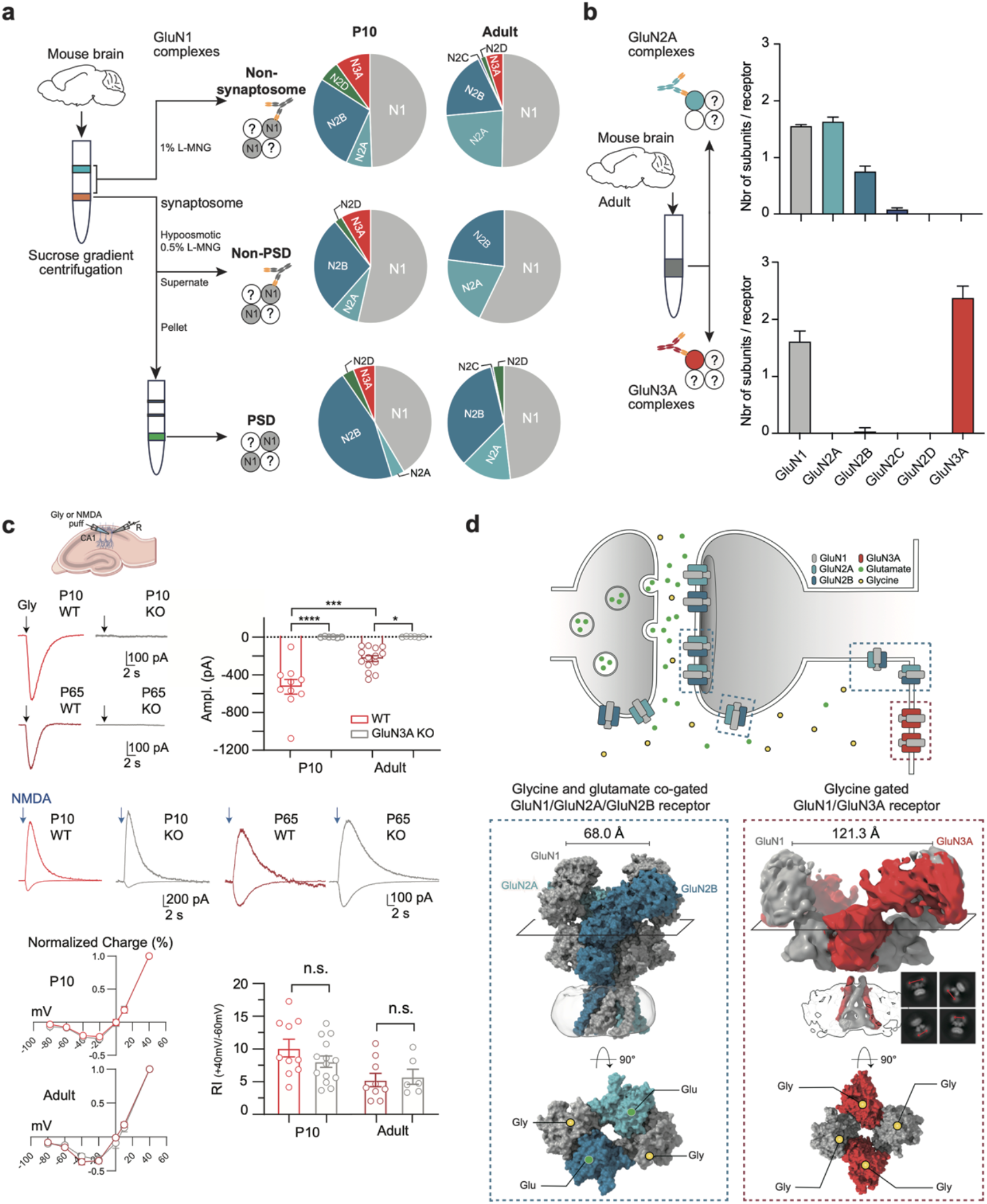
Developmental distribution, assembly, function, and structure of native GluN3A-containing NMDARs. (**a**) Mass spectrometric analysis of NMDAR subunit distribution in postnatal day 10 (P10) and adult mouse brain. Isolation of non-synaptosomal, non-PSD (postsynaptic density), and PSD fractions from brain homogenates following fractionation by sucrose density gradient centrifugation followed for the two formers by anti-GluN1 immunoprecipitation. The pie chart represents the relative abundance of NMDAR subunits in each fraction as determined by mass spectrometry. (**b**) Quantitative assessment of GluN subunits associated to GluN2A and GluN3A-containing NMDARs determined by targeted AP-MS from membrane fractions of adult mouse brains. Bar graphs illustrate molecular abundance values normalized to NMDAR tetramers for the indicated GluN subunits in GluN2A-(upper panel) and GluN3A-(lower panel) containing NMDARs. Data are mean (± SD) of three experiments.(**c**) Slice electrophysiological recordings of Glycine and NMDA puff on CA1 pyramidal neurons acute ventral hippocampal (VH). In top left traces and top right graph glycine puffs (1 mM) in CGP-78608 (1 μM) do induce currents (-65 mV) in WT but not in GluN3A KO juvenile (P10) and adult (> P65) mice. In the bottom part representative traces (-60, +40 mV), current-voltage relationship (I-V) and rectification index (Rl = I_AMP_ (+40) I_AMP_ (-60)) of currents elicited by pressure-applied NMDA (1 mM) are unchanged between WT and GluN3A-KO juvenile (P10) and adult (> P65) mice. (**d**) Model of an adult glutamatergic synapse showing the localization of glycine/glutamate-gated conventional GluN1/GluN2 receptors and glycine-gated eGlyRs (GluN1/GluN3A receptors). Bottom: Comparison of the corresponding NMDAR complexes: surface representations of GluN1/GluN2A/GluN2B receptors (PDB: 8XLK)^10^ and cryo-EM density map of wild-type eGlyRs from this study (Extended Data Fig. 2C). The transparent density corresponds to the detergent belt lying around the transmembrane region. The lower panels show top-down views of the ligand-binding domains (LBDs) with their corresponding agonists: glutamate bound to GluN2, and glycine bound to GluN1 and GluN3A subunits.

In contrast to canonical GluN2 subunits, the GluN3A subunit displayed distinct subcellular distribution and developmental shift patterns (Fig. 1a). At P10, GluN3A constituted a significant portion of the total NMDAR pool, estimated to 10-20%, and was detectable in all three fractions. The presence of GluN3A in P10 mouse synapse likely supports its transient and developmentally regulated roles in synapse maturation^20,23,24,43^. In the adult mouse brain, GluN3A was virtually absent from synaptosome compartments, but remained robustly expressed in the non-synaptosome fraction (Fig. 1a). This unique localization points to the distinct distribution and signaling of GluN3A-containing NMDARs.

### Native GluN3A-containing receptors assemble as glycine-gated GluN1/GluN3A diheteromers

We next assessed the subunit composition of native GluN3A receptors from mouse brain. We combined affinity-purifications via anti-GluN3A antibody on plasma-membrane enriched fractions with label-free quantitative mass spectrometry (AP-MS, see Methods). Similar procedures were performed on GluN2A affinity purified complexes. For GluN2A-containing receptors, these experiments demonstrate robust co-assembly with GluN1, GluN2B and GluN2C, in line with the formation of diheteromeric GluN1/GluN2A and triheteromeric GluN1/GluN2A/GluN2B and GluN1/GluN2A/GluN2C receptor complexes (Fig. 1b). This mixture is consistent with previously described subpopulations of native GluN2A-containing NMDARs^2,10,40,44^. In contrast, for GluN3A-containing NMDARs, our AP-MS experiments uncovered exclusive co-assembly of GluN3A with GluN1 at a stoichiometry of close to 2 for both subunits in the tetrameric complex (Fig. 1b). These data show that native GluN3A subunits are preferentially, if not exclusively, integrated into GluN1/GluN3A diheteromers (with two copies of GluN1 and two copies of GluN3A subunits).

We investigated the presence of functional GluN3A-receptors using electrophysiological recordings from acute ventral hippocampal slices, a region displaying strong GluN3A expression^30,31,45^. Puffing glycine on CA1 pyramidal cells in the presence of a cocktail of inhibitors (including the GluN1 antagonist CGP-78608; see Methods) resulted in large inward (i.e. excitatory) currents (Fig. 1c). Currents were absent in GluN3A KO mice, confirming the involvement of excitatory glycine GluN1/GluN3A receptors (eGlyRs). Currents were also larger in juvenile (P10) than adult animals, in agreement with GluN3A expression peaking during the first and second postnatal weeks^20,21,45^. To evaluate the presence of GluN1/GluN2/GluN3A triheteromeric receptors, current-voltage (I-V) relationships of puff-evoked NMDA responses were compared between WT and GluN3A KO slices^24,26,27,31,46^. No difference was observed in either P10 or adult animals (Fig. 1c). Therefore, biochemical and functional evidence indicates that in both the juvenile and adult brain, GluN3A-NMDARs assemble specifically as excitatory glycine-gated GluN1/GluN3A receptors (eGlyRs).

### Unique arrangement of glycine-bound GluN1/GluN3A receptors

We launched structural investigation of eGlyRs using a GluN1/GluN3A^EM^ receptor construct with the C-terminal domain (CTD) truncated to enhance protein yield and thermostability^5,6,47^. The GluN1/GluN3A^EM^ construct retained typical functional properties of wild-type (WT) eGlyRs, including profound glycine-induced desensitization and massive current potentiation by the GluN1 ligand CGP-78608^25,28^ (Extended Data Fig. 1a). Receptors were expressed, purified and supplemented with 1 mM glycine before being frozen on EM grids. 2D class averages of GluN1/GluN3A^EM^ receptors revealed diffuse densities of the extracellular domains (ECDs), indicative of conformational heterogeneity (Extended Data Fig. 2a-c). Final 3D refinement yielded a cryo-EM density map of the GluN1/GluN3A^EM^ receptor at a low resolution (∼8 Å, Extended Data Fig. 2c, Table 1) with distinguishable TMD, LBD and NTD layers (Fig. 1d). The density of GluN1/GluN3A^EM^ receptor matches an alternate GluN1/N3A/N1/N3A heterotetrameric organization but with loose packing of the ECD layers and separated NTD dimers, as if the subunits were splayed. A similar structural observation has been recently reported^48^.

This organization appears in stark contrast with the typical compact organization of GluN1/GluN2 NMDARs (Fig. 1d)^5,6,11,47^. Within the LBD layer, GluN1/GluN2 receptors adopt a typical two-fold symmetrical dimer-of-dimer organization, while in GluN1/GluN3A receptors, the GluN3A LBDs show a large reorganization, resulting in a pseudo four-fold rosette arrangement (Fig. 1d, lower panels).

In order to stabilize eGlyR structure and potentially improve structure resolution, we built a chimeric GluN1/GluN3A-(2B NTD) receptor, by replacing the NTD and NTD-LBD linker of the GluN3A subunits with the homologous region from GluN2B. The chimeric GluN1/GluN3A-(2B NTD) receptor retains WT eGlyR functional properties including profound desensitization and CGP-78608 potentiation (Extended Data Fig. 1b,d). Following purification and cryo-EM data collection (Extended Data Fig. 2d,e), 2D class averages of the GluN1/GluN3A-(2B NTD)^EM^ tetramer in the presence of glycine and CGP-78608 gave rise again to a density map with poor overall resolution (∼7Å, Extended Data Fig. 2e, Table 1). By masking the poor density of the NTD layer, we improved the resolution of the gating core region (LBD + TMD) to approximately 5.3 Å (Extended Data Fig. 2e) with an electron density clearly fitting CGP-78608 in the GluN1-LBD. The topology of the LBD layer again revealed rotated LBDs arranged in a pseudo four-fold symmetric manner (see Fig. 2e). Therefore, the overall shape of WT-like (GluN1/GluN3A^EM^) and chimeric (GluN1/GluN3A-[2B NTD]^EM^) eGlyR structures appears unique, diverging from all known GluN1/GluN2 NMDAR architectures (Fig. 1d). This divergence points to intrinsic structural differences determining the distinct functional properties of glycine-gated eGlyRs *vs* glycine/glutamate co-gated conventional NMDARs.

### Gain-of-function mutants at the LBD dimer interface

The D1-D1 dimer interface between adjacent LBDs is a critical and shared structural element controlling iGluR gating^1^. In our low-resolution glycine-bound GluN1/GluN3A^EM^ receptor structure, the two GluN3A LBDs undergo a conspicuous ∼103° counterclockwise rotation, compared to the configuration of GluN2 LBDs in conventional NMDARs (Fig. 2a). This large subunit reorientation has no equivalent in GluN2-receptors but resembles the LBD arrangement of certain AMPA and kainate receptors captured in desensitized states (see extended data Fig. 10)^49–52^. We thus hypothesized that the stability of the LBD dimer interface is lower in GluN1/GluN3A receptors than in GluN1/GluN2 receptors that exhibit only limited desensitization^1,53^. Interestingly, the residue distribution at the LBD dimer interface markedly differs between the two receptor types (Fig. 2a). In particular, a hydrophobic patch present in GluN2 LBD upper lobe (D1) is missing in GluN3A (grey circles, Fig. 2a). To investigate the role of this interface in eGlyR gating, we systematically substituted GluN3A residues by their corresponding GluN2 counterparts and tested the activity of mutated eGlyRs using electrophysiology.

**Figure 2.**
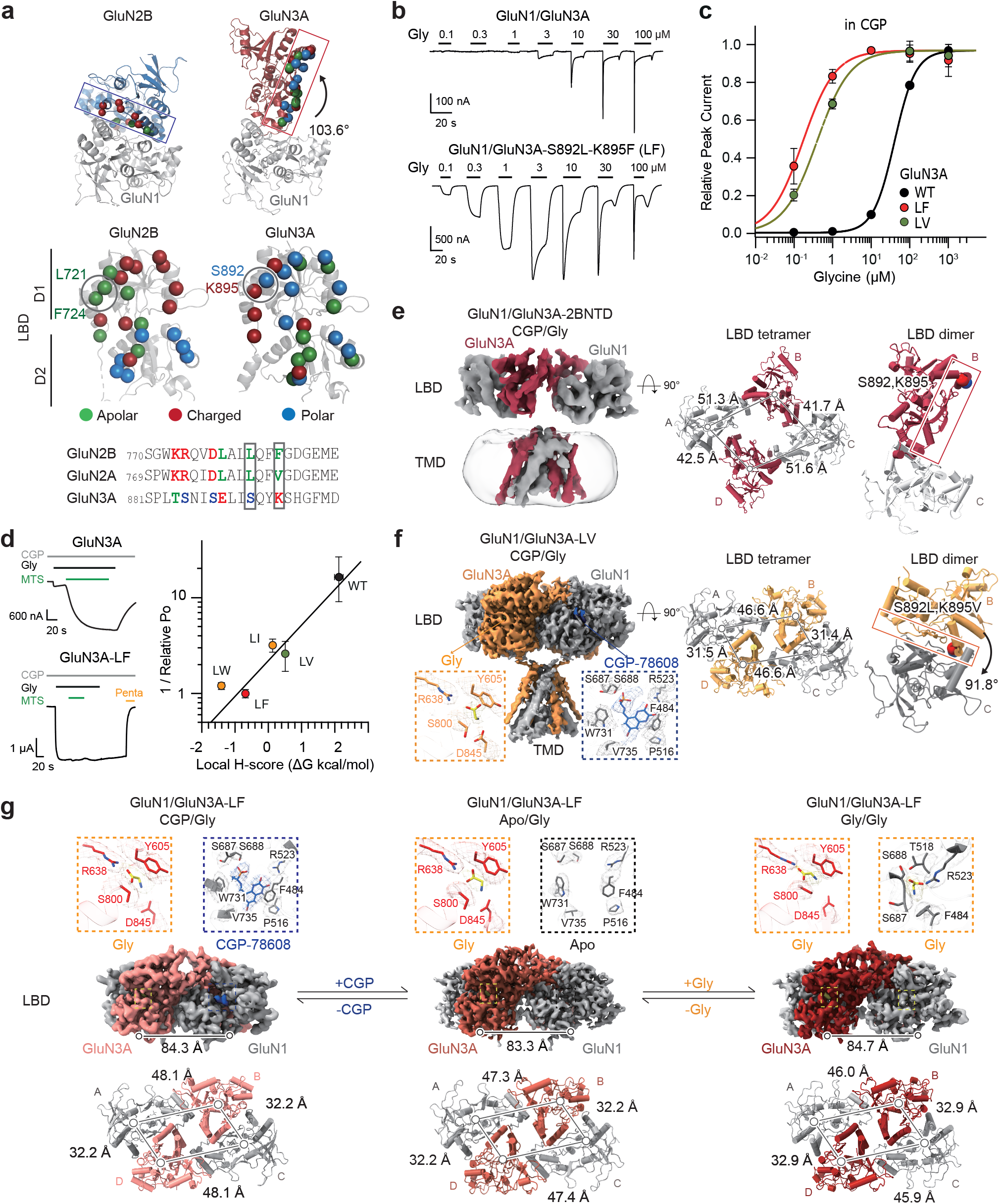
Structural and functional characterization of GoF1 mutants at the LBD dimer interface. (**a**) Top: LBD heterodimers in GluN1/GluN2B (PDB: 4TLL)^6^ and our GluN1/GluN3A-WT receptors structure. The rectangle marks the region corresponding to the known intra-LBD dimer contact interface. Middle: side-views of GluN2B-LBD and GluN3A-LBD. Spheres indicate the Cα atoms of residues of GluN2B and GluN3A that contact GluN1 at his interface. They are colored according to side chain properties: polar (blue), charged (red), and apolar (green). The gray circle highlights the position of the gain-of-function mutants GoF1. Bottom: sequence alignment of GluN3A, GluN2B, and GluN2A around the mutant sites. The amino-acids mutated in GoF1 are framed, with GluN3A-S892L/K895F (termed as GluN3A-LF) copying GluN2B while GluN3A-S892L/K895V (termed as GluN3A-LV) mimicks GluN2A. (**b**) Representative TEVC recordings from GluN1/GluN3A-WT and GluN1/GluN3A-LF receptors activated by glycine alone at different concentrations. (**c**) Glycine dose-response curves of WT and GoF1 receptors recorded in presence of 200 nM CGP-78608 to abolish desensitization. Error bars on points represent mean ± SEM. See Table 2 for values of EC_50_, Hill slope (n_H_) and statistics. See Extended Data Fig. 3 for complementary data and analysis. (**d**) GoF1 increase of relative open probability (Po). Left: Representative traces showing the effect of 300 µM MTSEA on GluN1/GluN3A-WT and GluN3A-LF mutant receptors together with GluN1-A652C mutant (see Methods). Glycine was applied at 100 µM and CGP-78608 at 200 nM. The constitutively active MTSEA-treated GoF1 mutants were inhibited by the pore blocker pentamidine (100 µM). Right: In the GoF1 background, relative 1/Po values of receptors with substitutions at GluN3A residues S892-K895 correlate with change in local hydrophobicity (H-score)^93^. See Table 3 for values of relative Po. Error bars represent mean ± SEM. (**e**,**f**) Cryo-EM structures of GluN1/GluN3A-2BNTD and GluN1/GluN3A-LV receptors. Structure determination: for panel E see Extended Data Fig. 2d,e and for panel F see Extended Data Fig. 4. Left: EM density maps of the LBD-TMD regions. Right: Top-down views of the LBD tetramer and dimer. Red spheres mark the Cα atoms of residues S892 and K895. Rectangles indicate the same interface than in (A). Rotation angles between intra-dimers were calculated using GluN1 as the reference. (F) Insets highlight the binding pockets for CGP-78608 and glycine in GluN1 and GluN3A, respectively. (**g**) Structures of GluN1/GluN3A-LF receptors trapped in CGP-78608/glycine, Apo/glycine, and glycine-bound states. For structure determination see Extended Data Fig. 5. Top: Cryo-EM density maps of the LBD regions, with insets showing binding pockets for glycine or CGP-78608. Bottom: Top-down views of the tetrameric LBD under the three conditions. The LBD-TMD linker distance between the two GluN3A subunits is indicated as a read-out of tension on the pore (see also Fig. 5). Distances between domains in panels e-g were measured between the center of mass (COM).

Among a series of mutants (Tables 2,3), we identified the double mutant GluN1/GluN3A-S892L-K895F (thereafter abbreviated as GluN1/GluN3A-LF) as a strong gain-of-function (GoF) mutant, dubbed GoF1 (Fig. 2a-d). GluN1/GluN3A-LF receptor responses differ from WT receptors by their enhanced current amplitude, enhanced sensitivity to glycine, and reduced desensitization (Fig. 2b-d, Extended Data Fig. 3a). The increase in glycine sensitivity is such (EC_50_ of 0.51 ± 0.03 µM [n=5] *vs* 8.1 ± 0.5 µM [n=5] for WT receptors) that we estimated GluN1/GluN3A-LF receptors to be partially (∼10%) activated by ambient glycine (Extended Data Fig. 3b) known to contaminate recording solutions (in the 10-50 nM range)^25^. We also obtained evidence that GluN1/GluN3A-LF receptors display greatly increased gating efficacy. While glycine-induced currents of WT GluN1/GluN3A receptors were massively (∼1000 fold) potentiated by CGP-78608^25^, GluN1/GluN3A-LF receptor currents showed only 2.4 ± 0.3-fold potentiation (n=12, Extended Data Fig. 3c, Table 3). This phenotype cannot be explained by a lower potency of CGP-78608 on the mutant receptors (CGP-78608 EC_50_ of 33.1 ± 1.2 nM [n=6] *vs* 26.3 ± 5 nM for WT receptors; Extended Data Fig. 3d)^25^. Rather, the strong reduction in the extent of CGP-78608 potentiation indicates high activity of GluN1/GluN3A-LF receptors. We obtained further supporting evidence by estimating the receptor’ maximal channel open probability (Po) using the MTSEA method (Fig. 2d). This method, developed on GluN1/GluN2 receptors, is based on the covalent modification of a cysteine residue introduced in the GluN1 subunit (GluN1-A652C), which locks open MTSEA cross-linked channels^54,55^. Accordingly, the extent to which the thiol-reactive MTSEA compound potentiates NMDAR currents is inversely correlated to the channel Po. WT receptors were potentiated 16.4 ± 7.3-fold (n=6) upon application of MTSEA (in the presence of 100 µM glycine and 200 nM CGP-78608, Fig. 2d), indicative of a low channel Po. In similar conditions, GluN1/GluN3A-LF receptors showed no potentiation by MTSEA (1.0 ± 0.1-fold potentiation [n=5]; Fig. 2d), indicative of a channel Po close to 1.0, the theoretical maximum (Table 3). Therefore, GoF1 double mutation at the LBD dimer interface (GluN3A-S892L-K895F) greatly increases receptor activity. Similar GoF effects were observed with the GluN3A-S892L-K895V mutations (GluN1/GluN3A-LV receptors; Fig 2c,d). Replacing GluN3A-K895 by various hydrophobic residues revealed a robust correlation between channel Po and hydrophobic volume at this site: the greater the hydrophobic volume, the higher the channel activity (Fig. 2d, Extended Data Fig. 3e and Table 3).

**Table 1.** Cryo-EM data collection, refinement and validation statistics for WT and Chimera eGlyRs.

|  | GluN1-4a/GluN3A<br>(Gly-Glu)<br>(EMD-62839;<br>PDB 9L5P) | GluN1-4a/GluN3A-(2B NTD)<br>(CGP-Glu)<br>(EMD-62851;<br>PDB 9L67) | GluN1-4a/GluN3A-(2B NTD)<br>Local refinement<br>(CGP-Glu)<br>(EMD-62850;<br>PDB 9L66) |
| --- | --- | --- | --- |
| **Data collection and processing** |  |  |  |
| Magnification | 105000 | 165000 | 165000 |
| Voltage (kV) | 300 | 300 | 300 |
| Electron exposure (e-/Å<sup>2</sup>) | 60 | 60 | 60 |
| Defocus range (µm) | -1 ~ -2 | -1 ~ -2 | -1 ~ -2 |
| Pixel size (Å) | 1.08 | 0.75 | 0.75 |
| Symmetry imposed | C2 | C2 | C2 |
| Initial particle images (no.) | 228,139 | 500,959 | 500,959 |
| Final particle images (no.) | 64,633 | 145,204 | 244,313 |
| Map resolution (Å) | 8.23 | 7.4 | 5.26 |
| FSC threshold | 0.143 | 0.143 | 0.143 |
| **Refinement** |  |  |  |
| Initial model used | SWISS MODEL | SWISS MODEL | SWISS MODEL |
| Model resolution (Å) | 8.07 | 7.36 | 5.66 |
| FSC threshold | 0.143 | 0.143 | 0.143 |
| Map sharpening B factor (Å<sup>2</sup>) | 495.00 | 450.00 | 190.00 |
| Model composition |  |  |  |
| Nonhydrogen atoms | 23658 | 25289 | 7155 |
| Protein residues | 2996 | 3196 | 898 |
| Ligands | 0 | 0 | 2 |
| B factors (Å<sup>2</sup>) |  |  |  |
| Protein | 585.0 | 510.30 | 372.85 |
| Ligand | --- | --- | 328.85 |
| Root-mean-square deviations |  |  |  |
| Bond lengths (Å) | 0.004 | 0.002 | 0.003 |
| Bond angles (°) | 1.088 | 0.619 | 0.811 |
| **Validation** |  |  |  |
| MolProbity score | 2.65 | 2.25 | 2.58 |
| Clashscore | 26.64 | 20.03 | 29.10 |
| Poor rotamers (%) | 2.36 | 0.18 | 0.00 |
| Ramachandran plot |  |  |  |
| Favored (%) | 92.80 | 92.82 | 86.69 |
| Allowed (%) | 6.86 | 7.15 | 13.20 |
| Disallowed (%) | 0.34 | 0.03 | 0.11 |

**Table 2.** Glycine EC_50_ and MgCl_2_ IC_50_ values for WT and mutant eGlyRs.

| Constructs | Glycine EC<sub>50</sub> |  |  | Glycine EC<sub>50</sub> in CGP-78608 |  |  |
| --- | --- | --- | --- | --- | --- | --- |
|  | peak current<br>(μM) | nH | n (cells) | (μM) | nH | n (cells) |
| GluN3A * |  |  |  |  |  |  |
| WT | 8.1 ± 0.5 | 1.28 | 5 | 45.9 ± 1.77 # | 1.59 | 5 |
|  |  |  |  | 43.6 ± 3.22 & | 1.75 | 11 |
| S892L-K895F (2x) | 0.51 ± 0.03 § | 1.3 | 5 | 0.16 ± 0.02 | 1.11 | 8 |
| S892L-K895V | 0.5 ± 0.11 | 1.4 | 5 | 0.39 ± 0.04 | 1.01 | 4 |
| S892L-K895I | 1.13 ± 0.62 | 1.38 | 5 | 0.64 ± 0.13 | 1.14 | 3 |
| S892L-K895W | N.M.<sup>1</sup> |  |  | 0.94 ± 0.13 | 1.05 | 4 |
| P497C | 48.5 ± 0.59 | 1.5 | 4 | 28.09 ± 0.65 | 2.01 | 5 |
| E498C | 45.0 ± 0.07 | 1.5 | 4 | 21.14 ± 2.85 | 1.79 | 4 |
| Q499C | 47.1 ± 1.00 | 1.5 | 4 | 28.75 ± 5.64 | 1.48 | 4 |
| A500C | 49.4 ± 3.01 | 1.5 | 4 | 21.92 ± 1.03 | 1.62 | 4 |
| Q501C | 35.5 ± 2.61 | 1.2 | 13 | 16.42 ± 0.67 | 1.95 | 10 |
| Q501A | N.M.<sup>2</sup> |  |  | 29.16 ± 1.23 | 2.05 | 7 |

\* : coexpressed with GluN1-4a ; #: dataset 1; &: dataset 2 ;
§ pour la composante d'inhibition : IC50 : 20.2 ; nH 1.25 ; max inhib 0.66

N.M.<sup>1</sup> Not measurable because of two strong tonic activation. N.M.<sup>2</sup> Not measurable because of too small current; N.D : Not determined.
| Constructs | MgCl<sub>2</sub> IC<sub>50</sub> |  |  |  |
| --- | --- | --- | --- | --- |
|  | IC50<br>(μM) | nH | Maximal<br>Inhibition | n (cells) |
| GluN3A-WT * | 3512 ± 237 | 0.94 | 1.00 | 4 |
| GluN3A-NN * (G729N-R730N) | 438 ± 70 | 1.1 | 0.66 | 4 |
| GluN3A-GN * (R730N) | 254 ± 25 | 1.5 | 0.46 | 5 |
| GluN2A-WT | 13.2 ± 2.6 | 1.0 | 0.99 | 3 |

\* : coexpressed with GluN1-4a

**Table 3.**
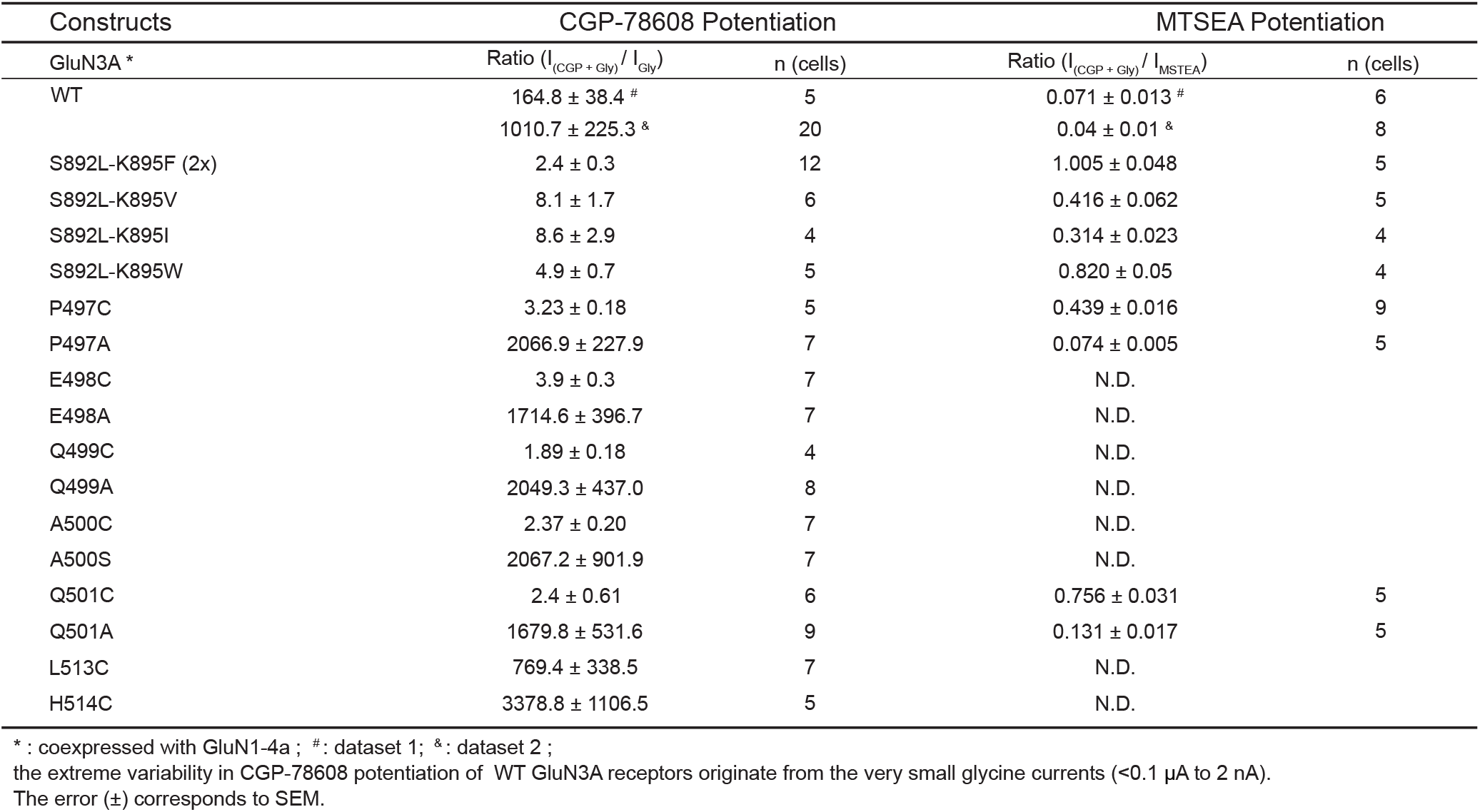
CGP-78608 and MTSEA potentiation in different eGlyRs.

| Constructs | CGP-78608 Potentiation |  | MTSEA Potentiation |  |
| --- | --- | --- | --- | --- |
| GluN3A * | Ratio ($I_{(CGP + Gly)} / I_{Gly}$) | n (cells) | Ratio ($I_{(CGP + Gly)} / I_{MTSEA}$) | n (cells) |
| WT | 164.8 ± 38.4 # | 5 | 0.071 ± 0.013 # | 6 |
|  | 1010.7 ± 225.3 & | 20 | 0.04 ± 0.01 & | 8 |
| S892L-K895F (2x) | 2.4 ± 0.3 | 12 | 1.005 ± 0.048 | 5 |
| S892L-K895V | 8.1 ± 1.7 | 6 | 0.416 ± 0.062 | 5 |
| S892L-K895I | 8.6 ± 2.9 | 4 | 0.314 ± 0.023 | 4 |
| S892L-K895W | 4.9 ± 0.7 | 5 | 0.820 ± 0.05 | 4 |
| P497C | 3.23 ± 0.18 | 5 | 0.439 ± 0.016 | 9 |
| P497A | 2066.9 ± 227.9 | 7 | 0.074 ± 0.005 | 5 |
| E498C | 3.9 ± 0.3 | 7 | N.D. |  |
| E498A | 1714.6 ± 396.7 | 7 | N.D. |  |
| Q499C | 1.89 ± 0.18 | 4 | N.D. |  |
| Q499A | 2049.3 ± 437.0 | 8 | N.D. |  |
| A500C | 2.37 ± 0.20 | 7 | N.D. |  |
| A500S | 2067.2 ± 901.9 | 7 | N.D. |  |
| Q501C | 2.4 ± 0.61 | 6 | 0.756 ± 0.031 | 5 |
| Q501A | 1679.8 ± 531.6 | 9 | 0.131 ± 0.017 | 5 |
| L513C | 769.4 ± 338.5 | 7 | N.D. |  |
| H514C | 3378.8 ± 1106.5 | 5 | N.D. |  |

\* : coexpressed with GluN1-4a ; #: dataset 1; &: dataset 2 ; the extreme variability in CGP-78608 potentiation of WT GluN3A receptors originate from the very small glycine currents (<0.1 µA to 2 nA). The error (±) corresponds to SEM.

### Structure of GoF1 receptors captured in an active-like state

We next investigated the structural correlates of the strong gain-of-function phenotype of GluN1/GluN3A LBD dimer mutant receptors. We purified the CTD-truncated GluN1/GluN3A-LV receptors (GluN1/GluN3A-LV^EM^) and GluN1/GluN3A-LF receptors (GluN1/GluN3A-LF^EM^) in the presence of glycine and CGP for single particle cryo-EM analysis (Extended Data Fig. 4-5). Receptor particles were collected under three different conditions: 1) in the presence of 1 mM glycine and 500 nM CGP-78608; 2) without supplementation of ligands; 3) with 1 mM glycine only. Through multiple rounds of 2D and 3D classification, we resolved cryo-EM structures of GluN1/GluN3A-LV^EM^ and GluN1/GluN3A-LF^EM^ receptors with local resolution of the gating core in the 3.1-3.5 Å range including well-defined densities of the LBDs (Fig. 2e-g, Extended Data Fig. 4-5 and Table 4). The four LBDs of GluN1/GluN3A-LV^EM^ receptors assemble as dimer-of-dimers with a pseudo two-fold symmetry and GluN1 subunits occupying the A-C positions (Fig. 2f), as typically observed in GluN1/GluN2 receptors^5,6,9,13^. Noticeably, the GluN3A-S892L-K895V mutations at the LBD heterodimer interface occupy the exact same positions as the corresponding residues in GluN1/GluN2 receptors (Fig. 2a). Similar results were observed with the GluN1/GluN3A-LF receptors (Extended Data Fig. 5). Thus, restoration of the hydrophobic D1-D1 inter-subunit contacts through the GluN3A double mutations appears sufficient to stabilize the LBD dimer interface in eGlyRs. Overall, the compact arrangement of the LBD layer in GluN1/GluN3A-LV and GluN1/GluN3A-LF receptors strikingly differs with the disrupted LBDs seen in WT-like eGlyRs (Fig. 1d) and GluN1/GluN3A-(2B NTD) receptors (Fig. 2e), indicating the capture of distinct, presumably active or pre-active, conformational states (see below).

In the three different ligand conditions, prominent densities assigned to orthosteric ligands are observed in the LBD central clefts. In the first condition, CGP-78608 and glycine occupy the cleft of GluN1 LBD and GluN3A LBD respectively (Fig. 2g), while in the presence of (saturating) glycine only, both GluN1 and GluN3A LBD harbor glycine molecules. In the condition without ligand added, a defined density for glycine is visible in the cleft of GluN3A LBD, but not in the GluN1 LBD (Fig. 2g). This aligns well with the known high glycine affinity of GluN3A^17^ and the constitutive activation of GluN1/GluN3A-LF receptors by low ambient glycine levels (see above). Therefore, we defined the three structures as CGP/Gly, Apo/Gly, and Gly/Gly bound states, respectively. The GluN3A LBD in complex with glycine adopted closed-cleft conformations in all three conditions (Fig. 2g, Extended Data Fig. 9). The GluN1 LBD also adopted a closed-cleft conformation in the Gly/Gly state, while in the absence of ligand (Apo/Gly) and in the presence of CGP-78608 (CGP/Gly), open-cleft conformations were observed, with angles augmented by 27.6° and 27.2°, respectively (Fig. 2g, Extended Data Fig. 9).

To get insights on the gating states of the GluN1/GluN3A-LF receptors, which display the strongest gain-of-function phenotype (Fig. 2d), we analyzed the inter B-D subunit distances (between α-helices E of the two GluN3A LBDs), a classic readout of the tension exerted by the LBD-TMD linkers on the channel pore^14,56^. In the three conditions, the measured distances were >80 Å, larger than in any other GluN1/GluN2 NMDARs described to date even for receptors captured in the pre-active or active state (Fig. 2g; see also Fig. 5a,b)^14,56–58^. These data indicate that GluN1/GluN3A-LF^EM^ receptor structures correspond to pre-active or active states in which the GluN3A LBD dimers exert strong pulling force on the ion channel pore. They also provide a structural correlate to the ability of eGlyRs to fully activate with only two agonist molecules bound (glycine on GluN3A subunits), a unique feature among iGluRs^25,29,34,35,59^.

### Large conformational mobility of the NTDs: crossed and uncrossed states

The poor resolution of the NTD layer of our imaged eGlyRs point to conformational heterogeneity of the NTDs. A similar conformational phenotype has been noted for GluA1-AMPA receptor complexes^50^. Detailed analysis in the initial rounds of 3D classification of both GluN1/GluN3A-LV andGluN1/GluN3A-LF receptors revealed at least two distinct conformations of the NTD layer (Extended Data Fig. 4 and 5), although the LBD layer remains in the active-like conformation described above. We termed these two conformations ‘crossed’ and ‘uncrossed’ states (Fig. 3a). In both states, the NTD layer arranges in a dimer-of-dimers configuration. In the crossed state, the two NTD dimers are closely interacting with each other through their GluN3A NTD R2 lobes, exhibiting a R2-R2 center-of-mass (COM) distance of 38.1 Å (Fig. 3a). In the uncrossed state, the two NTD dimers are positioned far apart (>60 Å; Fig. 3a), resembling the overall arrangement seen in the WT-like and chimeric eGlyR structures (Fig. 1d). Both crossed and uncrossed states exhibit loose NTD dimer assembly and poor interlayer compaction (Fig. 3a). Glued through R1-R1 interactions, individual NTD dimers appear widely extended, with the R2 lobes of GluN1 and GluN3A positioned almost orthogonally. This is in stark contrast with the arrangement observed in all other NTD dimers of iGluRs including GluN2-containing NMDARs (Extended Data Fig. 7a)^1,3,60^. This unusual NTD arrangement in eGlyRs creates substantial cavities between the NTD and LBD layers, with minimal interactions between the two layers.

**Figure 3.**
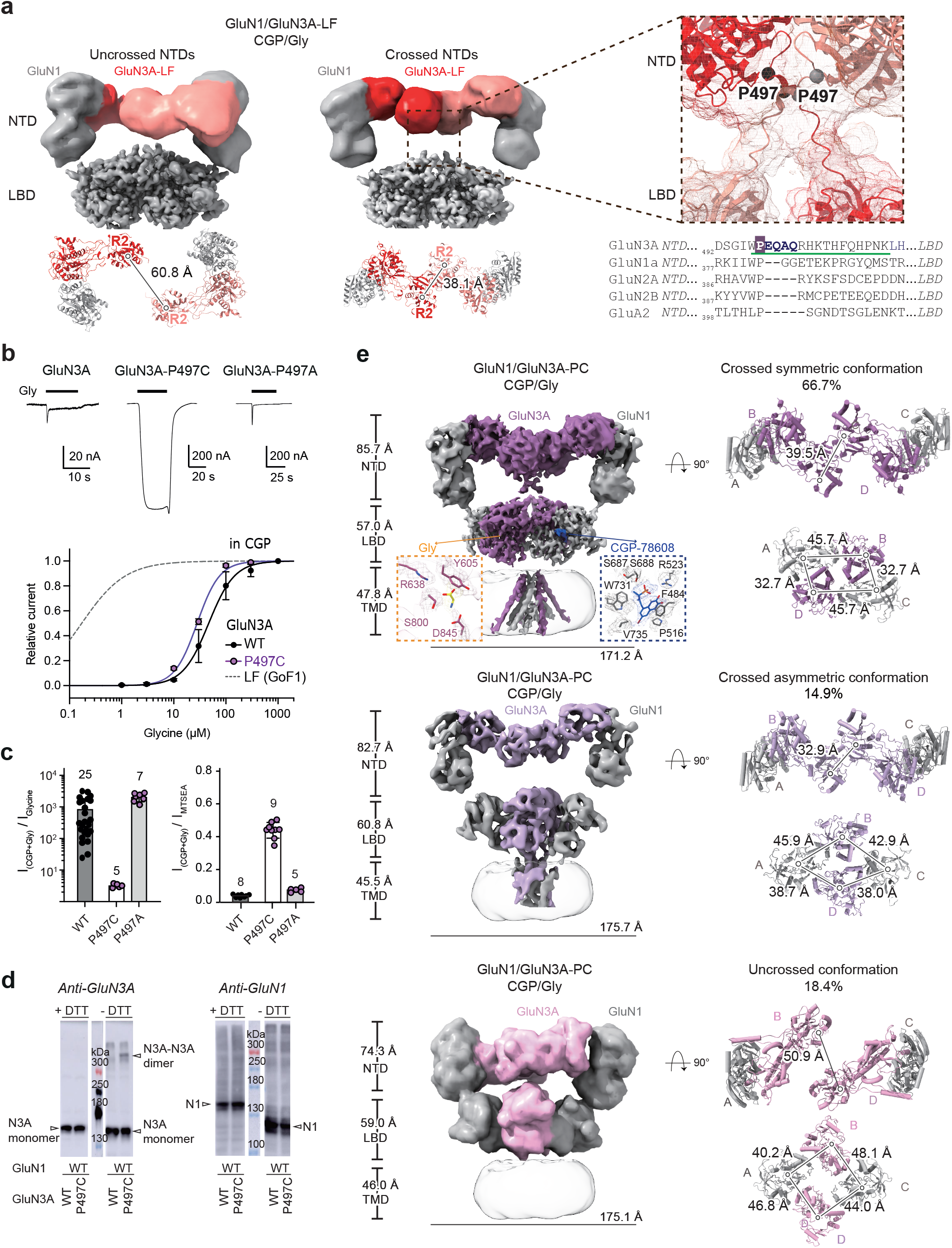
Structural and functional characterization of GoF2 crosslinking mutants in the GluN3A NTD-LBD linker. (**a**) Cryo-EM density maps of the extracellular region of GluN1/GluN3A-LF in the CCP-78608/glycine-bound state. Top-down views of the NTDs illustrate uncrossed (left) and crossed (middle) conformations, with GluN3A subunits colored red and orange (See Methods and Extended Data Fig. 7 for detailed analysis of NTD mobility in this receptors). Separation between NTD dimers was quantified as the COM distance of the R2 domains. On the right : zoom in the NTD-LBD linker region with the location of the GoF2 mutation (P487C or GluN3A-PC). Sequence alignment of the corresponding linker in other iGluRs receptors at the bottom. The green line position indicates the region corresponding to the linker, the tested mutant positions are in bold, while GoF2 position is highlighted in dark blue. (**b**) TEVC recording traces show responses to 100 µM glycine alone in GluN1/GluN3A, and control GluN1/GluN3A-PA receptors. Bottom: Glycine dose-response curves in the presence of 200 nM CGP-78608 for GluN3A-PC is compared to GluN3A-WT, GluN3A-LF (GoF1). Error bars on points represent mean ± SEM. See Table 2 for EC_50_ values and statistics. (**c**) GluN3A-PC shows drastic reduction of CGP-78608-induced potentiation compared to GluN3A-WT and PA, here quantified as the ratio of peak current in the presence of glycine + CGP-78608 to that with glycine alone. MTSEA-based measurements reveal a large increase of relative open probability (Po) in GluN3A-PC compared to control receptors. Error bars represent mean ± SEM. See Table 3 for values and statistics. (**d**) Immunoblots from HEK293 cells expressing WT and GluN1/GluN3A-PC receptors in either reducing and non-reducing conditions (DTT) and revealed with anti-GluN3A and anti-GluN1 antibody. Note the presence of a GluN3A-GluN3A dimer band in PC (n = 3 independent experiments). (**e**) Three distinct Cryo-EM structures of GoF2 (GluN1/GluN3A-P497C) receptors in CGP and Glycine. Left: Density maps of GoF2 receptors in crossed symmetric, crossed asymmetric, and uncrossed conformations, with insets highlighting densities at the agonist-binding sites. Right: Top-down views of the NTD and LBD layers in different conformations, with lines connecting the COMs of individual domains. The proportion of each conformation within the dataset is indicated in percentage.

We obtained further indication of the high conformational dynamics of eGlyR NTDs by performing *in silico* analysis of the potential transitions between the crossed and uncrossed states. For that purpose, we used iModFit^61^, an approach based on normal mode analysis that has proven useful in predicting large conformational changes of biomolecules including NMDARs^8,62,63^ (see Methods for details). The simulation, performed on (CTD-deleted) full-length receptors, revealed NTDs rearranging from the crossed to the uncrossed conformation following a progressive separation up to a full rupture of the inter-NTD dimer interface (Extended data Fig. 7b). The trajectory passes through structures present in our experimental datasets, yet not used as inputs in the fitting (Extended data Fig. 7c-e), supporting the realistic nature of the trajectory and high mobility of eGlyR NTDs.

### Locking the NTDs in the crossed state prevents eGlyR desensitization

Our structural and *in silico* observations of large mobility of eGlyR NTDs prompted us to explore the existence of such mobility in membrane-embedded receptors and its impact on receptor activity. Our structural data suggest that the crossed-NTD conformation might be a distinctive feature of active receptors, as it is observed only with the GoF1 receptors, while the uncrossed-NTD conformation with dissociated NTDs also exist in the desensitized-like state (Fig. 3a, Extended data Fig. 7a). We thus reasoned that stabilizing the crossed-NTD state should enhance receptor activity. To test this hypothesis, we introduced cysteine substitutions in the NTD-LBD linkers of GluN3A to restrict mobility following disulfide bridge formation (residues in bold; alignments of Fig. 3a). Our GoF1 cryo-EM structures indicate that in the crossed state, the two linkers from the two adjacent GluN3A subunits are sufficiently in close proximity while they are far apart in the uncrossed state (Fig. 3a). The five cysteine mutants tested (Fig. 3a, Extended Data Fig. 6 and Tables 2-3) including mutant GluN3A-P497C (GluN3A-PC) most proximal to the NTDs, showed particularly strong gain-of-function phenotype, which we named GoF2 mutants. In these mutants, application of saturating glycine (100 µM, no CGP-78608) triggered large and poorly desensitizing responses, in striking contrast to WT and alanine-substituted control receptors (Fig. 3b). Full glycine dose-response curves showed that, compared to GoF1 mutants, GoF2 mutants displayed only minor increase in glycine sensitivity but greatly reduced extent of desensitization even at high glycine concentrations (compare Fig. 2b and 3b; Table 2-3). As GoF1 mutants, the GoF2 mutants exhibited modest peak current potentiation by CGP-78608 (3.23 ± 0.18-fold [n=5] for GluN3A-P497C, and 2.4 ± 0.6-fold [n=6] for GluN3A-Q501C) compared to WT and alanine control receptors which were massively potentiated (∼1000 fold; Fig. 3c, Extended Data Fig. 6a and Table 3). Channel Po estimates confirmed that GoF2 mutant receptors displayed greatly enhanced channel activity (Fig. 3c, Table 3). Redox treatment experiments as well as western blot analysis of GoF2 mutant receptors confirmed the formation of disulfide bonds between the two GluN3A subunits (Fig 3d, Extended Data Fig. 6e,f). As additional control experiments, we introduced cysteine residues further downstream in the NTD-LBD linker, at positions GluN3A-L513C and GluN3A-H514C, which according to our structural data are too distant for crosslinking (>30 Å). Both GluN1/GluN3A-L513C and GluN1/GluN3A-H514C receptors responded to glycine as WT receptors (Extended Data Fig. 6a, Table 3).

### Intrinsic mobility of eGlyR gating core

We next leveraged GoF2 mutants to constrain the high mobility of NTD, and resolved the cryo-EM structure of the GluN1/GluN3A-PC receptor in the presence of glycine and CGP-78608 (Extended Data Fig. 8). Following multiple rounds of 2D and 3D classification, we identified three distinct conformational classes (Fig. 3e and Extended Data Fig. 8b): an NTD crossed conformation with LBDs arranged in a symmetric dimer-of-dimers, an NTD crossed conformation with the LBDs asymmetrically arranged with one dimer and two separated monomers, and an NTD uncrossed conformation with disrupted LBDs. The two NTD crossed conformations displayed GluN3A R2–R2 COM distances of 39.5 Å and 32.9 Å, respectively, while the uncrossed conformation had a distance of 50.9 Å (Fig. 3e). The symmetric conformation accounted for 66.7% of the particles and yielded the highest-resolution reconstruction. Applying a mask to exclude the NTDs improved the resolution of the gating core (LBD plus TMD) to 3.7 Å, allowing clear visualization of density for both glycine and CGP-78608 in the LBD clefts (Fig. 3e, Extended Data Fig. 8b).

Although we introduced the GluN3A-P497C as the GoF2 mutation into the NTD-LBD linker region, the gating core which contained no mutation showed distinct conformational states. It is striking that we can directly visualize the conformational flexibility of the LBDs within a single dataset, suggesting this mobility is likely part of the gating transition in WT eGlyRs. In the NTD crossed symmetric class, the COM distances within and between LBD dimers (32.7 Å and 45.7 Å, respectively) closely match those captured in the GoF1 structures (Fig. 2f-g). In contrast, in the NTD crossed asymmetric class, one dimer is broken, with the COM distance between GluN1-LBD and GluN3A-LBD increased to 42.9 Å, indicating an intermediate gating state captured by cryo-EM (Fig. 3e). The NTD uncrossed class comprises only 18.4% of the particles and yielded a reconstruction at relatively low resolution with disrupted LBDs. This class likely corresponds to a subpopulation of receptors lacking the disulfide bond formation at the GluN3A-P497C position since the inter-NTD distance is incompatible with such covalent bond. This uncrossed NTD conformation resembles those observed in WT eGlyR and GluN1/GluN3A-2B(NTD) structures (Fig. 1d, Extended Data Fig. 2). Notably, the crossed classes exhibit an elevated NTD layer, creating a large cavity between the NTD and LBD layers, a feature previously observed in most non-NMDA iGluRs^64^ (Fig. 3e). Taken together, these results establish that the crossed conformation observed in our cryo-EM data is a physiologically relevant structure of GluN1/GluN3A receptors. During the gating transition, the intrinsic mobility of the LBDs allows them to inter-convert from a disrupted tetramer into a reformed dimer-of-dimers configuration. Our data further demonstrate that the conformation of the NTD layer strongly influences eGlyR activity. Trapping the NTDs in the crossed conformation hinders entry into a desensitized state locking the receptor in a high Po mode.

### Structure and function comparison between conventional NMDARs and eGlyRs

The full-length structural elucidation of the GluN1/GluN3A-P497C receptor enables a direct structural comparison between conventional GluN2-NMDARs and glycine-gated eGlyRs. For this comparison, we used the glycine/glutamate-bound GluN1/GluN2B receptor (PDB: 7SAA^65^) and our glycine/CGP-78608-bound eGlyR structures, as both are trapped in agonist-bound, pre-active states. Globally, the eGlyR exhibits greater height (193.9 Å vs 170.1 Å) and width (172.2 Å vs 142.8 Å) than the GluN1/GluN2B receptor (Fig. 4a). This size difference primarily originates from the NTD layer. In eGlyRs, the GluN1/GluN3A NTD heterodimer is elevated, with the NTD-LBD linkers suspended between the two layers. This architecture provides both NTDs and LBDs with substantial space for motion, which ultimately influences channel activity as our GoF1 and GoF2 mutations. In contrast, in the GluN1/GluN2B receptor, the NTD layer sits directly atop the LBD layer, both layers acting as an integrated allosteric unit where NTD motion and regulation are primarily governed by the GluN2A and GluN2B subunits^8,10,11,55,57,62,66^.

**Figure 4.**
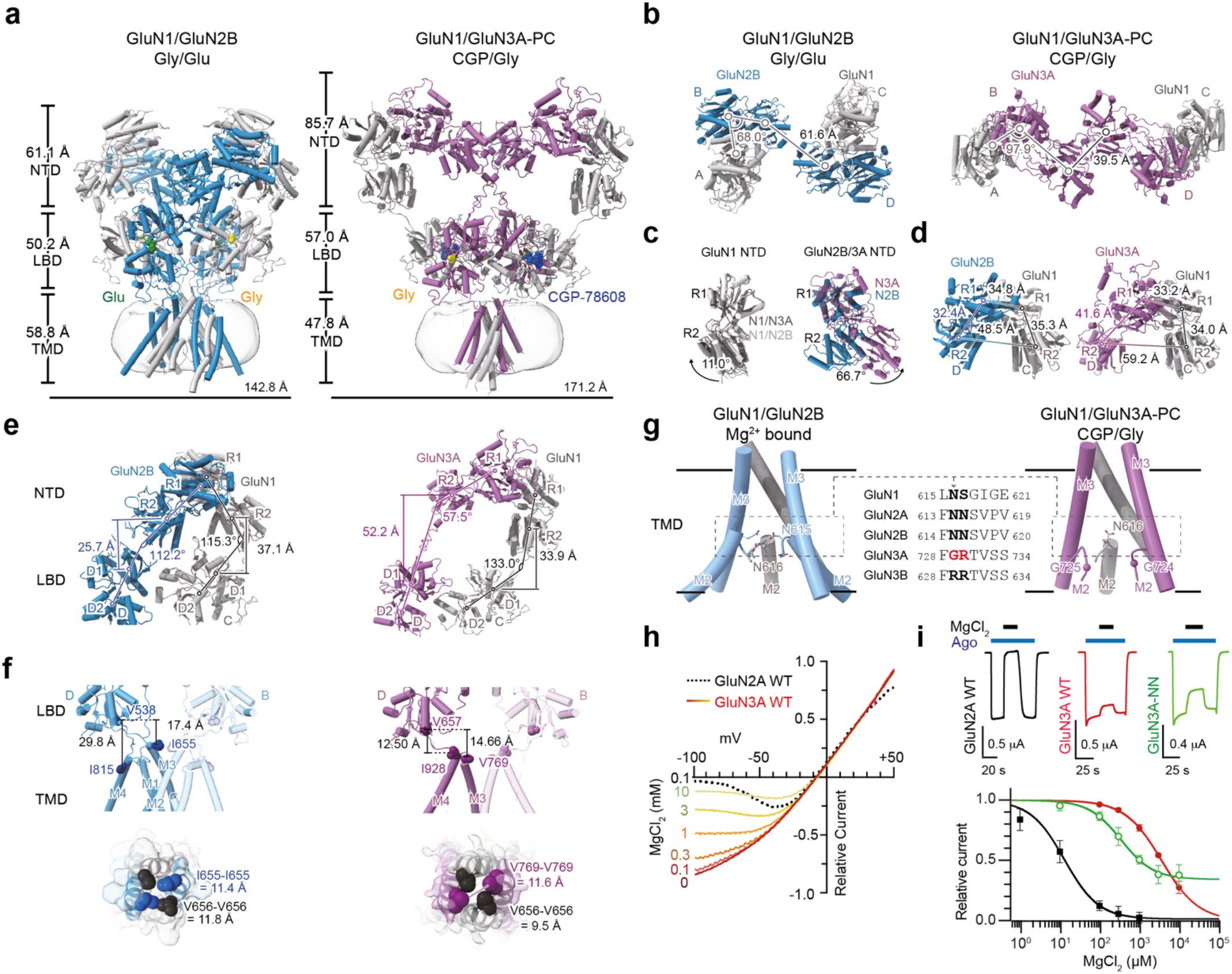
Structural comparison of GluN1/GluN2B and GluN1/GluN3A NMDA receptors. (**a**) Cartoon structural representations of the GluN1/GluN2B receptor (PDB: 7SAA)^65^ in the glycine/glutamate-bound state and the GluN1/GluN3A-PC receptor in the CGP-78608/glycine-bound symmetric state. (**b-d**) Structural comparison of top-down views for tetrameric NTDs (B), monomeric NTD (C), and heterodimeric NTDs (D) between GluN1/GluN2B and GluN1/GluN3A receptors. Domains are connected by lines between their centers of mass (COMs). For structural alignment in panel C, the R1 lobes were superimposed. (**e**) Structural arrangements between NTD and LBD layers in GluN1/GluN2B and GluN1/GluN3A receptors. Conformational differences between subunits are quantified by vector angles connecting the COMs of the R2 and R1 lobes of the NTDs, and the D1 and D2 lobes of the LBDs. (**f**) Comparison of the gating core (LBD plus TMD) between GluN1/GluN2B and GluN1/GluN3A receptors. Bottom panels highlight gate residues as spheres, with their Cɑ-Cɑ distances indicated. (**g**) Side view of the TMD in GluN1/GluN2B and GluN1/GluN3A-P497C receptor. The GluN1 subunit on the front has been removed for clarity. The dashed box indicates the QRN site location. In the middle is shown a sequence alignment of the TM2-TM3 segments of some human glutamate receptor subunits, with the QRN-site and N+1 site residues in bold. (**h**) Superimposed normalized I/V curves (recorded by TEVC in xenopus oocytes) of GluN1/GluN2 (dotted line) at 100 mM MgCl_2_ and GluN1/GluN3A-WT at various MgCl_2_ concentrations. (**i**) Top: TEVC traces of GluN2-WT, GluN3A-WT and GluN3A-NN (for GluN3A-G729N-R730N) recorded in the presence of saturating agonists and 300 mM MgCl_2_. Bottom: Corresponding MgCl_2_ dose-response curves for the three receptors. Error bars on points represent mean ± SEM. See Table 2 for IC_50_ and statistics.

At the level of individual domains, the GluN3A NTD adopts a more open cleft conformation, characterized by an R1-R2 distance of 41.5 Å. Additionally, its R2 lobes are rotated by 66.7° relative to those in the GluN2B NTD (Fig. 4b,c). By superimposing the R1 lobes, we observed that the R2 lobe of the GluN1 NTD undergoes an outward rotation of 11.0° in the eGlyR compared to its position in the GluN1/GluN2B receptor. Within the NTD heterodimer, the distance between the R2 lobes of GluN1 and GluN3A is also greater than that observed in the GluN1/GluN2B heterodimer (59.2 Å vs 48.5 Å, Fig. 4d). Furthermore, the interdomain angle between the NTD and LBD of the GluN1 subunit is significantly smaller in the eGlyR than in the GluN1/GluN2B receptor (47° *vs* 65.7°, Fig. 4e), a difference likely attributable to the outward displacement of the GluN1 NTD. Regarding the gating core transition between the LBDs and the TMD, the distance between these domains in the GluN3A subunit is considerably shorter than that in the GluN2B subunit (Fig. 4f). This structural feature may be linked to the fast-desensitization property characteristic of eGlyRs. At the ion channel gate, the key gate-forming residues in the eGlyR are arranged with two-fold symmetry, in contrast to the four-fold symmetry observed in GluN1/GluN2B receptors^12^. This distinct symmetry likely correlates with the fact that glycine binding to GluN1 or GluN3A triggers opposite gating effects in eGlyRs^25,29,34,35^.

The TMD pore domain of eGlyR displays a large vestibule lined by the four M3 helices above a constriction point formed by the M2 region (Fig. 4f,g), as typically observed in conventional NMDARs^1,12^. Although the resolution was insufficient to unambiguously assign side chains in the M2 region, we surmised that the GluN3A residues glycine 729 (G729) and arginine 730 (R730) at the tip of M2 (so-called Q/R/N site^1^), participate to the relative insensitivity of eGlyRs to Mg^2+^ block, another distinguishing feature with conventional NMDARs^26,33^. Voltage-ramps performed at various Mg^2+^ concentrations confirmed that eGlyRs are far less sensitive to Mg^2+^ block than GluN2A-containing NMDARs (IC_50_ value at -60 mV of 3512 ± 237 µM (n=4) *vs* 13.2 ± 2.6 µM (n=3), respectively; Fig. 4h,i). Substituting the GR pair in GluN3A with asparagines (NN), as present in GluN2 subunits, resulted in enhanced Mg^2+^ block of GluN1/GluN3A responses recorded at physiological membrane potential (Fig. 4i). These results identify the two GluN3A-specific M2 residues, G729 and R730, as distinctive molecular attributes contributing to the poor sensitivity of eGlyR channels to Mg^2+^ block.

## Discussion

In this study, we provide key information about the structure, mechanisms and molecular physiology of GluN3A-receptors, a subclass of NMDARs that has remained enigmatic for long. First, by employing quantitative mass spectrometry approaches on native brain extracts, we clarify long-standing issues about the abundance, subcellular distribution and assembly of GluN3A-receptors. We demonstrate that native GluN3A-receptors assemble as diheteromeric GluN1/GluN3A complexes that rely exclusively on glycine, and not glutamate, for their activation. We also show that GluN3A subunits undergo a major developmental regulation, being progressively excluded from synapses to be enriched at extrasynaptic compartments. Second, by combining single-particle cryo-EM, structure-guided mutagenesis and electrophysiological recordings, we decrypted the structural mechanisms underlying GluN1/GluN3A receptors unique gating and permeation properties. In particular, we identify key regions of the receptor accounting for their profound desensitization and resistance to Mg^2+^ block, distinguishing properties with conventional GluN2-NMDARs. Overall, our work establishes a robust molecular, cellular and physiological foundation to the emerging field of GluN3A-mediated excitatory glycinergic signaling and its translational potential. It has long been thought that native GluN3A subunits assemble with GluN1 and GluN2 subunits to form GluN1/GluN2/GluN3A triheteromeric receptors, co-activated by glutamate and glycine as conventional GluN2-NMDARs^67^. However, this model was recently challenged by the observation that GluN1/GluN3A diheteromeric receptors, initially discovered in recombinant systems^33^, are functionally expressed in several brain regions where they retain their insensitivity to glutamate and activation by glycine alone^25–32^. The relative abundance of the two receptor subtypes remained controversial^20,21^. Our current results based on mass spectrometry and electrophysiology establish that native GluN3A subunits assemble as glycine-gated GluN1/GluN3A diheteromers (i.e. eGlyRs) and not as GluN1/GluN2/GluN3A triheteromers (Fig. 1). Even in the juvenile brain, where GluN3A expression is highest^20^, our experimental evidence points to GluN3A subunits incorporating into eGlyRs. Therefore, native GluN1, GluN2, and GluN3 subunits follow an exclusion rule that yields separate populations of GluN1/GluN2 and GluN1/GluN3 diheteromers, echoing previous results obtained on recombinant NMDARs^68,69^. The recognition that endogenous GluN3A subunits assemble as GluN1/GluN3A eGlyRs prompts to revisit the many roles of GluN3A previously attributed to GluN1/GluN2/GluN3A receptors^67^. The eGlyRs being insensitive to glutamate (no GluN2 subunits), it also shed new light on glycine neurotransmission in the brain.

Our mass spectrometry data also establish that in the juvenile brain (P10), GluN3A subunits are the 2^nd^ most abundant non-GluN1 subunits (behind GluN2B), pointing to widespread roles of eGlyRs and glycine signaling in brain development. Our data further inform that at early postnatal ages GluN3A subunits are broadly distributed on the neuronal surface, including at synaptic sites, but undergo a major redistribution in adulthood being excluded from synaptic sites. Combined with our functional data, we therefore propose that in the adult brain eGlyRs localize extrasynaptically where they act as glycine sensors regulating neuronal excitability (volumic glycinergic signaling^27,31^). In contrast to conventional GluN2-NMDARs, GluN3A-containing eGlyRs lack PDZ-binding motifs, likely accounting for their loose anchoring at synaptic sites and high membrane diffusivity^69,70^. We also propose that in the developing brain, a significant fraction of eGlyRs is present at synapses, and mediate the described roles of GluN3A-receptors in synapse maturation^20,23,24,43^. How juvenile eGlyRs are activated, and whether these receptors operate at the post-or pre-synapse^67,71^ remains to be uncovered.

Our identification of strong GoF1 and GoF2 mutants, together with cryo-EM structures of WT and mutant eGlyRs resolved in distinct conformational states under various ligand conditions (glycine and/or CGP-78608), enabled us to propose a comprehensive model of eGlyR gating mechanisms (Fig. 5). In its fast-desensitizing WT form (Fig. 1d and 2e), eGlyR predominantly adopts a conformation characterized by disrupted tetrameric LBD architecture. This is structurally evidenced by the connections between the COMs of individual LBDs forming a pseudo-square shape, loose contact between GluN1 and GluN3A with an interfacial surface area of 327.7 Å^2^, and a short distance of approximately 26.9 Å between the α-helix E regions of two GluN3A LBDs (Fig. 5a,b; right). This latter structural parameter typically reflects the tension applied to pull open the ion channel gate^1^. The short distance is interpreted as weak tension, and under this condition, the gate remains trapped in the closed and desensitized state of WT eGlyR.

**Figure 5.**
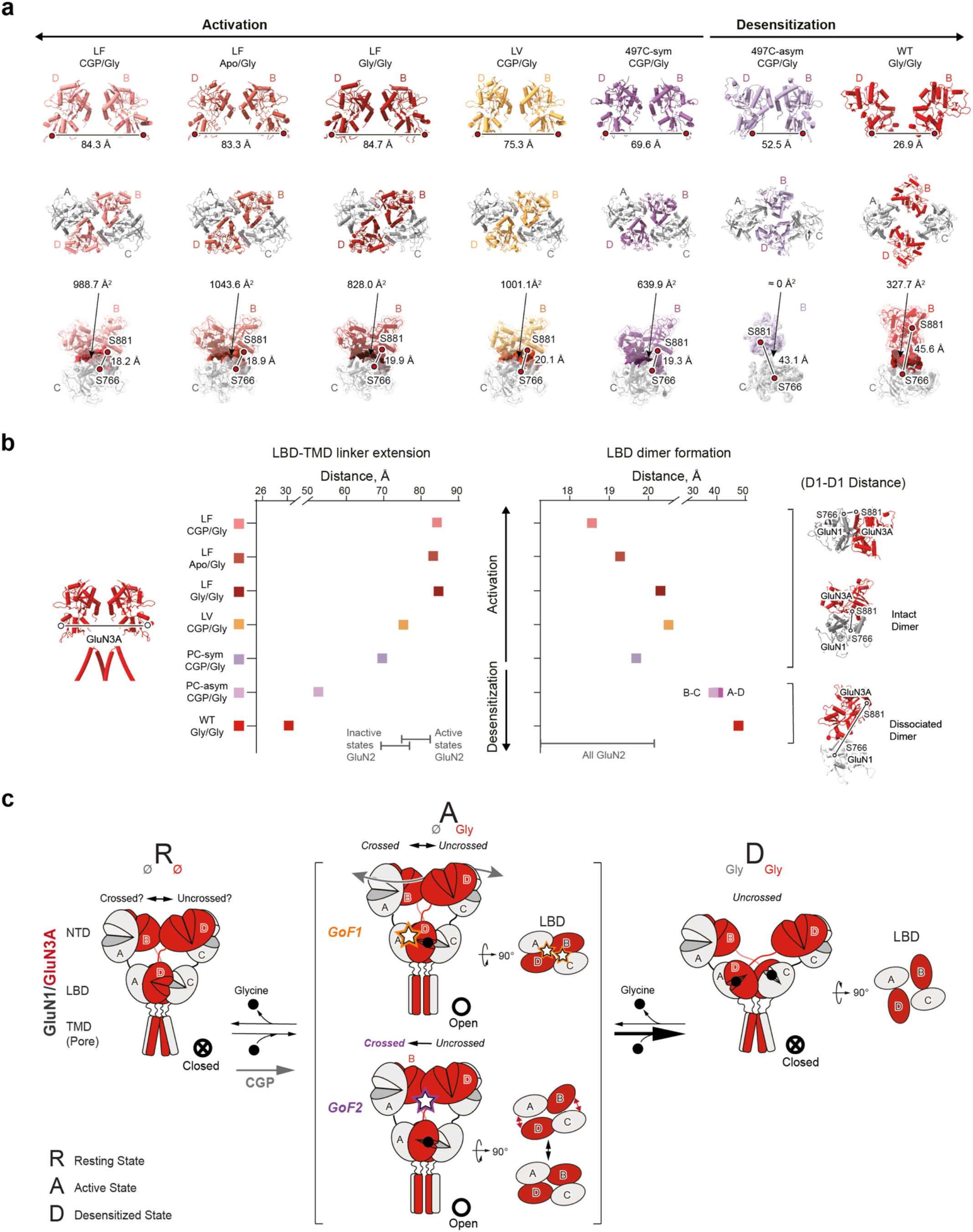
Model of eGlyR gating mechanism. (**a**) Representation of important gating metrics for our structural sample of active to desensitized states. Top line: Side-view cartoon representation of the two GluN3A LBDs for each of our eGlyR structures with the measured distances between the α-helix E regions of in all our structures. This LBD-TMD linker extension measurements are used as a proxy for the tension exerted by the LBDs on the transmembrane pore^65^. Middle line: Structural comparison of the LBD tetramer organization of each structure in top-down view. Bottom line: zoom in LBD dimers with intra-dimer D1-D1 distance measurement and contact surface size (non-transparent surface) between GluN1 and GluN3A. This former is used as a proxy for LBD dimer dissociation. (**b**) Graphical representation for the values of the LBD-TMD linker extension (left) and that of the intra-dimer LBD distance (right). For additional information see also Extended Data Fig. 9. The range of values measured in GluN2 structures are shown for comparison, revealing that LBD-dimer interface is already distended in LF Gly/Gly and LV CGP/Gly compared to the stable GluN2 dimers. (**c**) Proposed model illustrating how conformational changes in the LBD and NTD layers drive the eGlyR gating mechanism from Resting to Active and to Desensitized state upon binding of glycine. GoF1 and GoF2 mutants stabilize differently the active state. GoF1 mutant acts as non-covalent glue at the LBD-dimer interface while covalent GoF2 mutants impair full disruption into four-fold LBD arrangement. GoF2 also reveals the existence of a partly desensitized eGlyR transition state where the LBDs are separated but didn’t undergo transition to four-fold state. On the other hand, the NTDs display substantial conformational mobility, fluctuating between crossed and uncrossed conformation, unless in the fully Desensitized state where only uncrossed conformations were observed.

Introduction of GoF1 (both LV and LF) mutations at the GluN3A LBD markedly enhanced channel Po (Fig. 2), indicating stabilization of active eGlyR states. Structurally, these mutations promote the formation of canonical heteromeric LBD dimers with substantially increased interfacial contact surface area (1001 Å^2^ in LV and >820 Å^2^ in LF) and considerably reduced COM distances between GluN1 LBD and GluN3A LBD (∼48 Å in WT *vs* ∼31 Å in LV and ∼32 Å in LF). Consequently, the distance between the α-helix E regions of the two GluN3A-LBDs is greatly expanded to 75.3 Å in LV and above 83 Å in LF (Fig. 5a,b; left). Such large distances are observed even on eGlyRs with glycine bound only to GluN3A subunits (Apo/Gly structure; Fig. 5a,b). This is in line with the fact that glycine binding to the sole GluN3A subunits directly triggers channel gate opening in eGlyRs^25,29,34,35^. The gating core architecture observed in these GoF1 mutants resembles the dimer-of-dimers conformation typically seen in conventional GluN2-containing NMDARs^5,6,10^.

Alternatively, introduction of the GoF2 (GluN3A-P497C) mutation at the NTD-LBD linker region of the GluN3A subunit elevated channel Po to ∼0.5 and markedly suppressed desensitization (Fig. 3). Structural elucidation of a panoramic view of the GoF2 mutation within the WT eGlyR gating core background revealed substantial LBD conformational mobility (Fig. 3e). Notably, in the predominant crossed symmetric structure of the GoF2 receptor, disulfide crosslinking of the linker region distally drives the formation of a dimer-of-dimers architecture in the LBDs, characterized by extensive heterodimeric interfacial contact of 639.9 Å^2^, a short COM distance of 32.7 Å, and an extended distance of 69.6 Å between the two α-helix E regions (Fig. 5a,b; middle). These observations suggest that the WT eGlyR background retains capacity to transition between desensitized state and active dimer-of-dimers configuration. An asymmetric conformation was also captured, with structural parameters intermediate between those of the WT and active states, likely representing an intermediate state along the gating transition pathway (Fig. 5a,b; middle).

Integrating all structures resolved in this study, we observed a clear conformational trend in the LBD-TMD linker extension: progressing from loose tension (short distance) in the desensitized WT eGlyR, to an intermediate state in the moderately high Po GoF2 mutants, and ultimately to a wide separation to pull the opening of the gate in the active GoF1 mutants (Fig. 5b, left panel). Conversely, the conformational trend in the LBD heterodimer formation follows the opposite pattern. The WT receptor exhibited the largest inter subunit separation, GoF2 mutants showed intermediate distances, and GoF1 mutants displayed the closest contact between GluN1 LBD and GluN3A LBD (Fig. 5b, right panel). Altogether, our work reveals distinct structural and gating features of glycine-gated eGlyRs (Fig. 5c): their active states resemble those of conventional glycine/glutamate co-gated GluN2-containing NMDARs, while their desensitized states exhibit LBD disruption reminiscent of certain AMPARs and kainate receptors (Extended Data Fig. 10)^49–52^. Another salient feature of eGlyRs revealed by our structural data is their uniquely elevated and extended NTDs, likely increasing the mobility of the LBDs (Fig. 4). This differs from the compact NTD-LBD arrangement observed in GluN2-NMDARs^5–11,57^. We also show high conformational mobility of eGlyR NTDs, as observed in GluA1-type AMPARs^50,60^, with splaying apart of the two NTD dimers accompanying eGyR transition to the desensitized state.

Overall, eGlyRs appear as hybrid iGluRs in terms of both structure and function. With AMPA and kainate receptors, they are prone to profound desensitization due to an unstable LBD dimer interface^50,51,72^. They are also capable of channel gating despite partial occupancy of agonist binding sites^73–76^. With GluN2-containing NMDARs, their closest paralogs, they share the necessity for glycine binding to activate^3,19^. However, with their specific M2 pore residues (Fig. 4), eGlyRs lack the typical high Mg^2+^ block of GluN2-NMDARs. We believe these properties are tailored to match the physiological attributes of eGlyRs: sensors of extracellular glycine mediating tonic excitatory currents that regulate neuronal excitability^21,27,31^. Finally, we foresee strong pharmacological potential for compounds stabilizing specific eGlyR conformations^77^. We predict the GluN1-GluN3A LBD heterodimer interface to be a potential locus for small molecule allosteric potentiators, as described in other iGluRs^1,78,79^. The unprecedented large cavities between the NTDs and LBDs also offer opportunities for drug targeting.

## RESOURCE AVAILABILITY

### Data and code availability

The atomic coordinates for eGlyR structures have been deposited in the PDB with following accession codes PDB 9L5P (for GluN1/GluN3A), 9L67 and 9L66 (for GluN1/GluN3A-(2B NTD)), 9L5S, 9L5N and 9L5M (for GluN1/GluN3A-LF), 23IS (for GluN1/GluN3A-LV), and 23JH, 23OZ and 230V (for GluN1/GluN3A-P497C); see Tables 1, 4 and 5. The corresponding EM maps have been deposited in the EMDB with accession codes: EMD-62839, EMD-62851, EMD-62850, EMDB-62827, EMDB-62838, EMDB-62837, EMDB-69002, EMDB-69016, EMDB-69137 and EMDB-69134, respectively. The paper does not report original code. All materials are available from the corresponding authors upon request.

**Table 4.** Cryo-EM data collection, refinement and validation statistics for GluN1/GluN3A-S892L,K895F and GluN1/GluN3A-S892L,K895V receptors.

|  | GluN1-4a/GluN3A-LF<br>(CGP/Gly)<br>(EMDB-64046;<br>PDB 9UCK) | GluN1-4a/GluN3A-LF<br>(APO/Gly)<br>(EMDB-62838;<br>PDB 9L5N) | GluN1-4a/GluN3A-LF<br>(Gly/Gly)<br>(EMDB-64045;<br>PDB 9UCJ) | GluN1-4a/GluN3A-LV<br>(CGP/Gly)<br>(EMDB-69002;<br>PDB 23IS) |
| --- | --- | --- | --- | --- |
| **Data collection and processing** |  |  |  |  |
| Magnification | 165000 | 165000 | 165000 | 129500 |
| Voltage (kV) | 300 | 300 | 300 | 300 |
| Electron exposure (e-/Å<sup>2</sup>) | 60 | 60 | 60 | 60 |
| Defocus range (µm) | -1~-2 | -1~-2 | -1~-2 | -1~-2 |
| Pixel size (Å) | 0.74 | 0.74 | 0.74 | 0.93 |
| Symmetry imposed | C2 | C2 | C2 | C2 |
| Initial particle images (no.) | 491,660 | 406,908 | 334,747 | 696,964 |
| Final particle images (no.) | 336,538 | 65,646 | 60,386 | 99,176 |
| Map resolution (Å) | 3.07 | 3.23 | 3.33 | 3.50 |
| FSC threshold | 0.143 | 0.143 | 0.143 | 0.143 |
| **Refinement** |  |  |  |  |
| Initial model used | SWISS MODEL | SWISS MODEL | SWISS MODEL | SWISS MODEL |
| Model resolution (Å) | 3.0 | 3.2 | 3.4 | 3.50 |
| FSC threshold | 0.143 | 0.143 | 0.143 | 0.143 |
| Map sharpening B factor (Å<sup>2</sup>) | 158.4 | 167.0 | 175.7 | 118.7 |
| Model composition |  |  |  |  |
| Nonhydrogen atoms | 9,042 | 10,008 | 8,978 | 10,721 |
| Protein residues | 1,138 | 1,326 | 1,388 | 1,428 |
| Ligands | 8 | 6 | 10 | 4 |
| B factors (Å<sup>2</sup>) |  |  |  |  |
| Protein | 32.22 | 40.74 | 89.01 | 64.14 |
| Ligand | 33.10 | 37.29 | 102.57 | 59.97 |
| Root-mean-square deviations |  |  |  |  |
| Bond lengths (Å) | 0.006 | 0.007 | 0.007 | 0.006 |
| Bond angles (°) | 0.624 | 0.631 | 0.714 | 0.803 |
| **Validation** |  |  |  |  |
| MolProbity score | 1.67 | 1.64 | 1.94 | 1.95 |
| Clashscore | 4.76 | 4.07 | 8.00 | 15.63 |
| Poor rotamers (%) | .0.00 | 0.40 | 0.32 | 0.22 |
| Ramachandran plot |  |  |  |  |
| Favored (%) | 93.56 | 93.11 | 91.44 | 96.26 |
| Allowed (%) | 6.44 | 6.89 | 8.47 | 3.52 |
| Disallowed (%) | 0.00 | 0.00 | 0.09 | 0.10 |

**Table 5.**
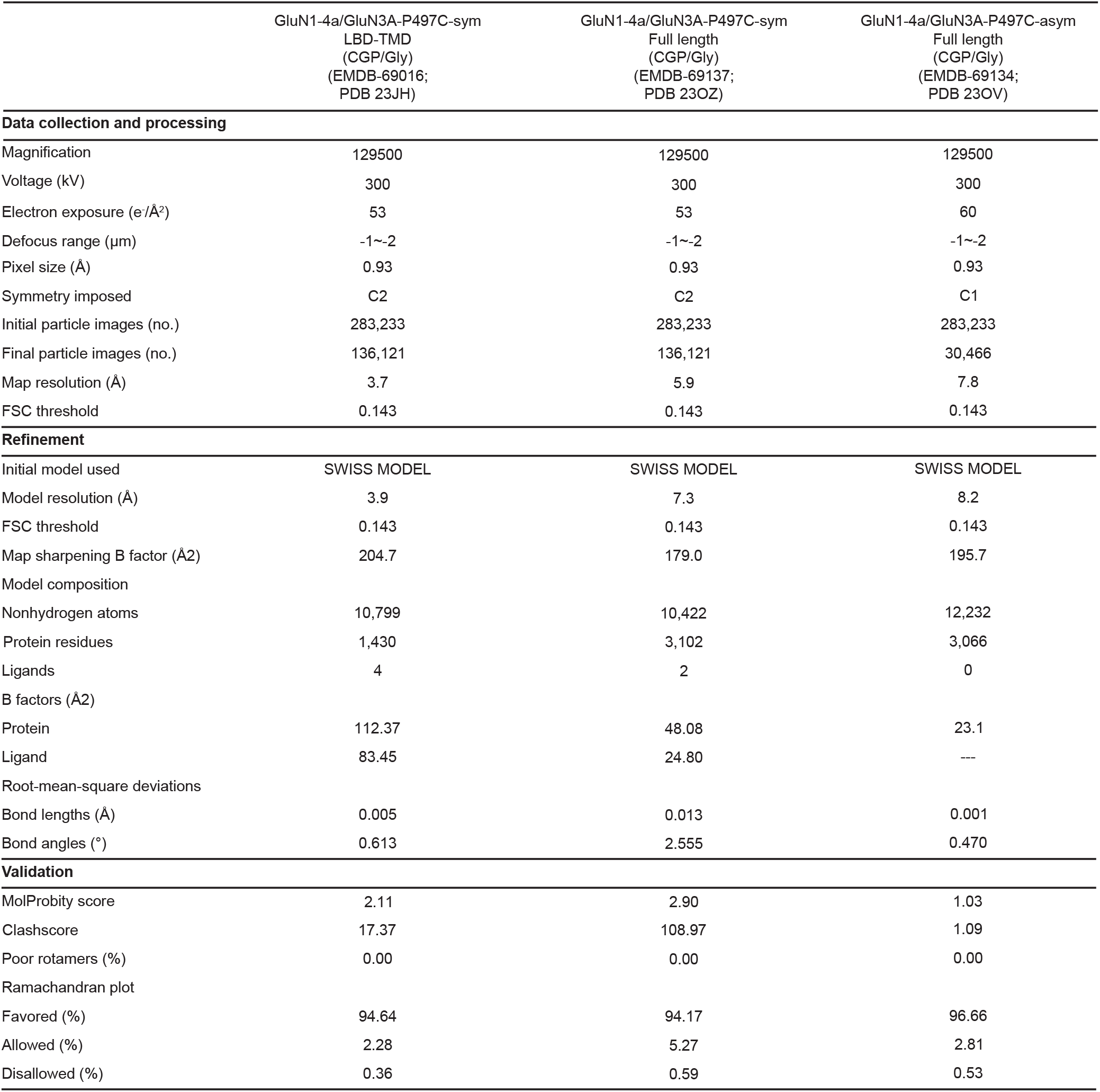
Cryo-EM data collection, refinement and validation statistics for GluN1/GluN3A-P497C receptors.

**Table 6.** Dataset of GluN1/GluN2A structures used for the structural analysis used in Fig. 5.

| PDB Code | Receptor Group | Technic/<br>reso (Å) | Conditions | Corresponding<br>construct | Reference |
| --- | --- | --- | --- | --- | --- |
| 7EOS | 2A Pre-Act | EM / 3.9 | Gly+Glu, pH8.0 | No CTD,cysteine mutations | Han Wang et al. 2021 |
| 7EOR | 2A Pre-Act | EM / 4.0 | Gly+Glu+GNE-6901, pH8.0 | No CTD,cysteine mutations | Han Wang et al. 2021 |
| 7EOQ | 2A Non-Act (Orthosteric inhibition) | EM / 4.1 | Gly+CPP, pH8.0 | No CTD,cysteine mutations | Han Wang et al. 2021 |
| 7EOT | 2A Non-Act (Orthosteric inhibition) | EM / 3.8 | CGP-78608+Glu, pH8.0 | No CTD,cysteine mutations | Han Wang et al. 2021 |
| 6IRA | 2A Pre-Act | EM / 4.5 | Glu+Gly, pH 7.8 | No CTD,multiple mutations | Zhang et al. 2018 |
| 6MMU | 2A Pre-Act | EM / 5.3 | Glu+Gly, pH 7.4 | No CTD | Jalali-Yazdi et al. 2018 |
| 6MMV | 2A Pre-Act | EM / 4.7 | Glu+Gly, pH 7.4 | No CTD (map with no TMD) | Jalali-Yazdi et al. 2018 |
| 6MML | 2A Pre-Act | EM / 7.1 | Glu+Gly, pH 7.4 | No CTD | Jalali-Yazdi et al. 2018 |
| 6MMS | 2A Pre-Act | EM / 5.4 | Glu+Gly, pH 7.4 | No CTD | Jalali-Yazdi et al. 2018 |
| 6MMR | 2A Pre-Act | EM / 5.1 | Glu+Gly, pH 7.4 | No CTD | Jalali-Yazdi et al. 2018 |
| 6MMG | 2A Pre-Act | EM / 6.2 | Glu+Gly, pH 7.4 | No CTD | Jalali-Yazdi et al. 2018 |
| 6MMW | 2A Pre-Act | EM / 6.2 | Glu+Gly, pH 7.4 | No CTD | Jalali-Yazdi et al. 2018 |
| 6MMA | 2A Non-Act (Allosteric inhibition) | EM / 6.3 | Glu+Gly, pH 6.1 | No CTD | Jalali-Yazdi et al. 2018 |
| 6MMX | 2A Non-Act (Allosteric inhibition) | EM / 7.0 | Glu+Gly, pH 7.4 | No CTD | Jalali-Yazdi et al. 2018 |
| 6MM9 | 2A Non-Act (Allosteric inhibition) | EM / 6.0 | Glu+Gly, pH 6.1 | No CTD | Jalali-Yazdi et al. 2018 |
| 6MMK | 2A Non-Act (Allosteric inhibition) | EM / 6.1 | Glu+Gly, pH 7.4 | No CTD | Jalali-Yazdi et al. 2018 |
| 6IRF | 2A Non-Act (Allosteric inhibition ) | EM / 5.1 | Glu+Gly, pH 6.3 | No CTD,multiple mutations | Zhang et al. 2018 |

**Table 7.**
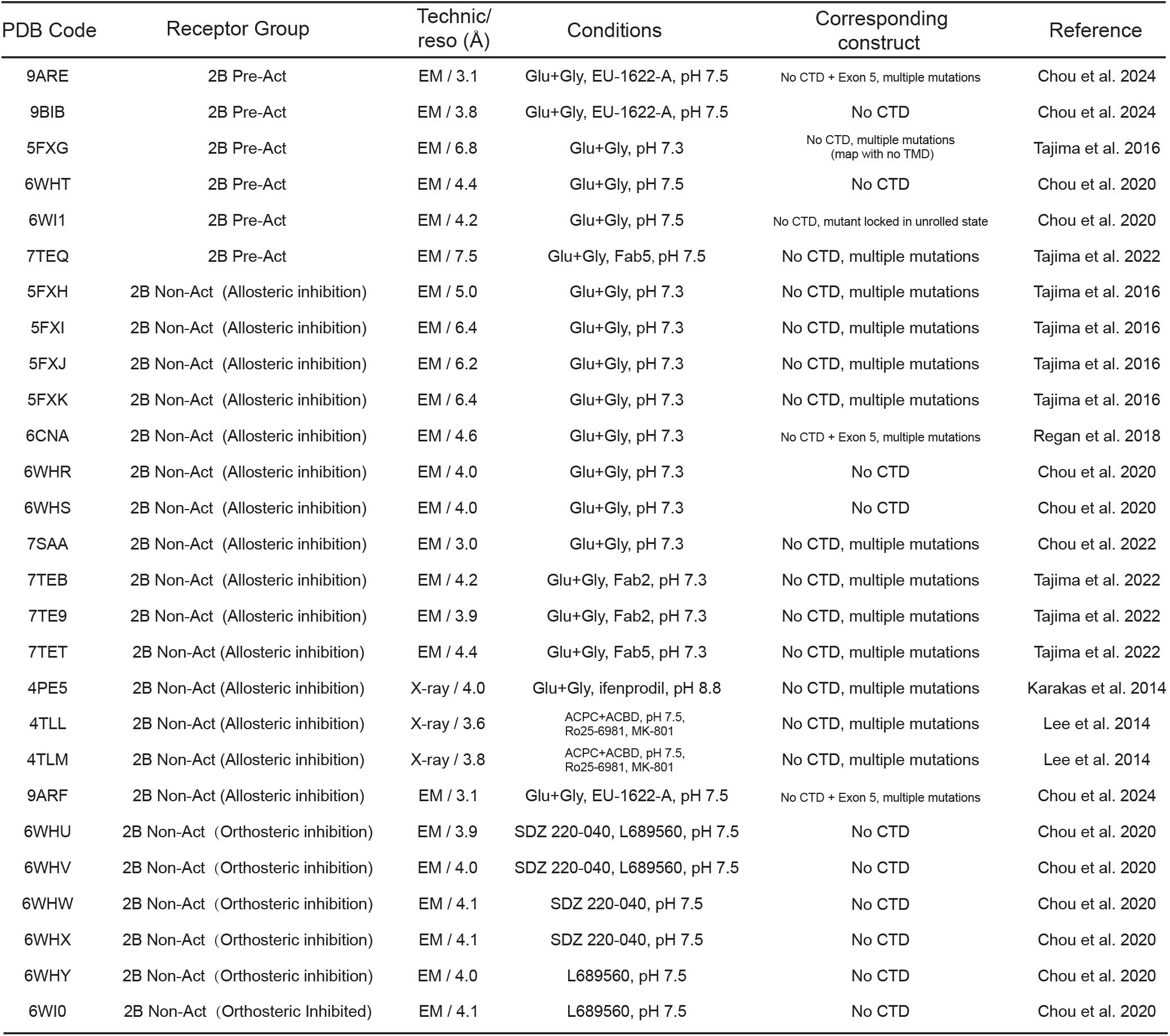
Dataset of GluN1/GluN2B structures used for the structural analysis used in Fig. 5.

## ACKNOWLEDGMENTS

This work was supported by the National Natural Science Foundation of China (32221003 to S.Z.), the SUSTech Distinguished Young Scientist Team Program and New Cornerstone Science Foundation through the XPLORER PRIZE (to S.Z.), the STl2030 Major Projects of China (2022ZD0212700 to S.Z.), the China Postdoctoral Science Foundation (2022M723247 to L.X.), the Agence Nationale de la Recherche (MEMOLIFE ANR-10-LABX-54, IDEX ANR-11-IDEX-0001-02, and ANR EXCIGLY to P.P.), the European Research Council (ERC Advanced Grant #693021 to P.P.), PSL University (Major Research Program ‘PSL-Neuro’, ANR-10-IDEX-0001 to P.P.), the Fondation pour la Recherche Médicale (FRM, Équipe FRM - EQU202503020034 to P.P.), the Région Ile-de-France (DIM C-BRAINS), the Deutsche Forschungsgemeinschaft (grants Fa 332/21-1 and ID 431984000 to B.F.), and Sorbonne Université (PhD fellowship to L.N.). We are grateful to the IBENS informatics facility for their technical support. We thank Simon Bossi, Siming Wu and Ke Xu for help with the preparations of mouse brains, and Cécile Cardoso for mouse genotyping.

## AUTHOR CONTRIBUTIONS

L.X. conducted all protein production, structural assays and cryo-EM data processing, HEK electrophysiology experiments and analysis under the guidance of S.Z. and D.S. L.X. analyzed all structural data. D.S., M.B.D. and L.N. designed GoF mutants. M.D.B., L.N. and A.C. carried out the TEVC and WB experiments and analysis under the guidance of D.S. and P.P. L.P. performed the electrophysiological experiments and analysis on brain slices. K.Y., J.S. and B.F. performed the native purification and mass spectrometry experiments and analysis. L.X. and D.S. assembled the figures of the manuscript. L.X., D.S., S.Z. and P.P. wrote the manuscript. D.S., S.Z. and P.P. supervised the study.

## METHODS

### Detailed Methods can be found in the supplementary file

#### Statistical Analysis

Unless otherwise indicated, data are presented as mean ± standard deviation of the mean (SEM). Non-parametric tests were used to assess statistical significance. When comparing two conditions, the non-parametric Mann Whitney test was employed, while for more than two groups the non-parametric Kruskal-Wallis test was employed, with Dunn’s multiple comparisons post hoc test. Statistical significances are indicated with *, **, *** when p-values are below 0.05, 0.01 and 0.001 respectively. n.s. indicates non-significant.

