## Supplementary_material_Xu_et_al for "Distribution, assembly and mechanism of GluN1/GluN3A excitatory glycine receptors"

#### METHODS

##### Mass Spectrometry on native brain extracts for determination of NMDAR subunit abundance and distribution

***-Purification of PSD, non-PSD and non-synaptosome fractions***

PSD, non-PSD and non-synaptosome fractions were isolated from the whole brain of C57BL/6J mouse (P10 and 8-week) as described previously (Suzuki, T. in *Neuroproteomics* (ed Ka Wan Li) 47–61, Humana Press, 2011). In brief, 5 grams of mouse brain were homogenized in solution A (0.32 M sucrose, 1 mM $MgCl_2$, 0.5 mM $CaCl_2$, 1 mM $NaHCO_3$) by 7 up-and-down strokes using a motor-operated homogenizer. After two-step low-speed centrifugations at 1,475 g and 755 g, for 10 min respectively, to remove nucleus, the supernatant (S1) was centrifuged at 17,300 g for 10 min and the pellet (P2) was collected. The supernatant (S2) was retained on ice. P2 was resuspended with solution B (0.32 M sucrose, 1 mM NaHCO3) and layered on gradients composed of 1.0/1.4 M sucrose, and centrifuged at 82,500 g for 70 mins. The synaptosome band in the interface between 1.0 and 1.4 M sucrose layers was collected. The upper layers were collected and combined with S2 as the non-synaptosome fraction. The synaptosome was added dropwise into 1 mM $NaHCO_3$ in beakers and stirred for 40 min at 4°C. And an equal volume of 1% L-MNG/0.32 M sucrose/12 mM Tris-HCl (pH 8.1) was slowly added while stirring. After stirred for 12 min, it was centrifuged at 48,2000 g for 20 min and the supernatant was collected as the non-PSD fraction. The pellet was homogenized in solution B and layered on gradients composed of 1.0/1.5/2.1 M sucrose, and centrifuged at 201,800 g for 120 mins. The band in the interface between 1.5 and 2.1 M sucrose layers was collected. It was diluted and centrifuge at 40,000 g for 20 min to get the PSD pellet for mass spectrometry analysis.

***-Affinity purification of NMDARs from non-synaptosome and non-PSD fractions***

To increase the abundance of NMDARs for mass spectrometry analysis, NMDARs were affinity-purified from detergent-solubilized membranes derived from non-synaptosome and non-PSD fractions. The non-synaptosome fraction was firstly centrifuged at 40,000 g for 40 min to get the membrane fraction. The pellet was then resuspended and solubilized in TBS with 1% L-MNG for 2 h at 4℃. After centrifugation at 40,000 g for 40 min, excessive GluN1 antibody Mab4F11 and protein G resin (Beyotime, SA016025) were added into the supernatant, and stirred for 2 h at 4℃. The resin was rinsed with wash buffer (TBS supplemented with 0.1% L-MNG) and collected by centrifugation at 4,000 g for 3 min. For the non-PSD NMDARs purification, excessive GluN1 antibody Mab4F11 and protein G resin were directly added into the non-PSD fraction. Finally, after stirred for 2 h at 4℃, the resin was rinsed and collected for mass spectrometry analysis.

***-LC-MS/MS analysis***

The PSD, non-synaptic and non-PSD NMDARs samples were firstly denatured by 8 M urea and subsequently followed the incubation of 10 mM dithiothreitol (DTT) at 55℃ for 25 min, 30 mM iodoacetamide (IAA) at RT for 30 min and 5 mM DTT at RT for 15 min in the dark. Samples were added with 7 volumes of 50 mM Tris-HCl (pH 8.5) and 1 mM $CaCl_2$, and digested with trypsin at 1:20 mass ratio at 37 ℃ for 14 hours. After added with 1% formic acid (FA), samples were centrifuged at 1,100 g for 10 min to remove the pellets. The supernates were loaded onto C18 StageTip columns (Thermo Fisher,

87784), which were activated by 80% acetonitrile (ACN) plus 0.5% acetic acid and equilibrated by 1% FA. After washed twice with 1% FA, samples were eluted with 80% ACN plus 0.5% acetic acid. The eluate peptides were dried under vacuum.

Mass spectrometry analysis was performed on an Orbitrap Fusion mass spectrometer (Thermo Fisher Scientific) equipped with a nanospray ion source and coupled to an EASY-nLC 1000 system (Thermo Fisher Scientific). Peptides were directly loaded onto an analytical column (100 μm inner diameter × 20 cm) packed with 1.9 μm, 120 Å ReproSil-Pur C18 resin. Peptides were separated using a binary mobile phase system consisting of buffer A (0.1% FA in water) and buffer B (0.1% FA in ACN). Elution was performed with a linear gradient of 7–22% buffer B over 100 min, followed by 22–35% buffer B over 20 min. Full MS scans were acquired in the Orbitrap over an m/z range of 350–1550 at a resolution of 120,000 with a target of $2 \times 10^5$ ions. MS/MS spectra were collected in data-dependent acquisition mode using top-speed method with higher-energy collision dissociation (HCD), with a target of $5 \times 10^4$ ions, a maximum injection time of 40 ms, an isolation window of 1.6 m/z, and a normalized collision energy of 30%. Dynamic exclusion was enabled with a duration of 30 s. Precursor ions with unassigned charge states, as well as those with charge states of 1+ or greater than 6+, were excluded from fragmentation.

***-Data Analysis***

Data from two independent mass spectrometry experiments were combined for analysis. The final abundance values for each of the six NMDAR subunits represented the mean value of two independent datasets. Distribution profiles across the PSD, non-PSD, and non-synaptosome fractions were visualized via pie charts processed in GraphPad Prism 9.

**Mass spectrometry on native brain extracts for determination of GluN subunit assembly and stoichiometry**

Membrane-enriched protein fractions were prepared from brains of adult and P8 mice, as described[80]. Dissected brains were homogenized in 15 mL HP-buffer (20 mM HEPES pH 7.5, 300 mM Sucrose, 1.5 mM $MgCl_2$, 1 mM EGTA, protease inhibitors (PMSF, Aprotinin, Leupeptin, Pepstatin A)) with a 15 ml Dounce homogenizer. Homogenates were centrifuged (10 min, 1,000g) and supernatants transferred into ultracentrifugation tubes. After ultracentrifugation (20 min at 200,000xg), the pellets were homogenized in 25 ml Lysis buffer (5 mM Tris/HCl pH 7.4 supplemented with protease inhibitors) and incubated on ice for 30 minutes. Additional ultracentrifugation (20 min at 200,000xg) separates soluble proteins from the membrane/membrane attached proteins. The latter was resuspended in 20 mM Tris/HCl pH 7.4 and loaded on top of a two-layer density gradient (0.5 M /1.3 M Sucrose in 10 mM Tris/HCl pH 7.4). After ultracentrifugation (45 min at 30,000 rpm, Beckman SW-30), the membrane interface was harvested and washed with 20 mM Tris/HCl pH 7.4. A final pellet after ultracentrifugation was resuspended in 20 mM Tris/HCl pH 7.4 and the protein concentrations were determined by Bradford assay and adjusted to 10 mg/ml. We pooled samples from mice of the same genotype (3x WT, 2x GluN3A KO).

For affinity purifications (APs), plasma-membrane enriched fractions were used as source material and specificity of APs was controlled by target-knockout material (GluN3A) and APs with preimmunization IgGs. For each AP, 1-1.5 mg of membrane proteins from WT or GluN3A KO mice were solubilized with 1-1.5 ml CL-47 (Logopharm) supplemented with protease inhibitors (Aprotinin, Leupeptin, Pepstatin A, PMSF). Homogenates were cleared by ultracentrifugation (10 min at 125,000xg) and solubilisates were incubated with 10-15 μg antibodies pre-coupled to protein A Dynabeads. We used the following antibodies: rabbit polyclonal anti-GluN3A (Sigma Aldrich, #07-356), rabbit monoclonal anti-GluN2A (Sigma Aldrich, #04-901), rabbit polyclonal anti-GluN2A (Sigma Aldrich, #07-632). Antibodies were incubated for two hours with solubilisates. As controls, we incubated anti-GluN3A antibodies with solubilisates from target KO, and IgGs (Millipore, #12-370) with WT solubilisates. Subsequently, beads were washed two times with 0.5 ml CL-47 buffer and proteins eluted with 10 μl of Lämmli buffer w/o DTT.

***-Mass spectrometry (MS) and protein quantification***

AP eluates were separated on SDS-Gel and in-gel digested with trypsin. The extracted peptide mixtures dissolved in 0.5% (v/v) trifluoroacetic acid were analyzed by nano-LC-MS/MS on an Orbitrap Elite (Thermo Fisher Scientific GmbH, Dreieich, Germany). Vacuum-dried peptides were dissolved in 20 µl of 0.5% (v/v) trifluoroacetic acid, loaded onto a trap column (C18 PepMap100, 5 µm particles, Thermo Fisher Scientific GmbH, Dreieich, Germany), separated by reversed-phase chromatography via a 10 cm C18 column (PicoTip™ Emitter, 75 µm, tip: 8 µm, New Objective, self-packed with ReproSil-Pur 120 ODS-3, 3 µm, Dr. Maisch HPLC; flow rate 300 nl/min) using an UltiMate 3000 RSLCnano HPLC system (Thermo Scientific), and eluted by an aqueous organic gradient (eluent "A": 0.5% acetic acid; eluent "B" 0.5% acetic acid in 80% acetonitrile. MS-analyses were executed on an Orbitrap Elite mass spectrometer with a Nanospray Flex Ion Source (both Thermo Scientific). Precursor signals (LC-MS) were acquired with a target value of 1,000,000 and a nominal resolution of 240,000 (FWHM) at m/z 400; scan range 370 to 1700 m/z). LC-MS/MS data were extracted using "msconvert.exe" (part of ProteoWizard; http://proteowizard.sourceforge.io/). Peak lists were searched against a UniProt Reference Proteome database (mouse) with peptide mass tolerance ± 5 ppm; fragment mass tolerance ± 0.8 Da using Mascot 2.6.2 (Matrix Science, UK). One missed trypsin cleavage and common variable modifications including S/T/Y phosphorylation were accepted for peptide identification. Significance threshold was set to $p < 0.05$. Abundancenorm values (as a measure of molecular abundance) were calculated as described[81]. Briefly, peptide signal intensities (peak volumes, PVs) were extracted from FT full scans and mass calibrated using MaxQuant v1.6.3 (http://www.maxquant.org). Peptide PV elution times were then aligned and assigned to peptides based on matching m/z and elution times (tolerances 2-3 ppm / ± 1 min). $\text{Abundance}_{\text{norm}}$ spec values (as a measure of molecular abundance) were calculated as described[81]. Determination of specificity in affinity purifications (APs) was based on target-normalized abundance ratios (tnRs) of proteins in WT versus control APs (calculated as described[82]). Normalized molecular abundance values for the specifically purified NMDA receptor pore forming subunits were obtained by relating the molecular abundance values of individual GluN proteins to the number of NMDA receptors via division by the factor ($\sum$ of GluN subunit abundances)/4.

#### Molecular biology

For two-electrode voltage-clamp electrophysiology in Xenopus oocytes, the rat GluN1 splice variant 4a (hereafter referred to as GluN1) construct was cloned into the pcDNA3 vector, while the rat GluN3A construct (UniProt ID: Q9R1M7) was cloned into the pcI_NEO vector, as described[25]. For whole-cell patch-clamp electrophysiology on HEK cells, full-length and CTD-truncated rat GluN1 and GluN3A constructs, as well as full-length human GluN1-1a and GluN2B constructs, were cloned into the pcDNA3.1 vector, with enhanced green fluorescent protein (eGFP) fused to the C-terminus of the GluN1 subunit. For recombinant protein expression for structural studies, CTD-truncated rat GluN1 and GluN3A constructs were cloned into a bicistronic pEG-BacMam vector. A 3C protease cleavage site (LEVLFQGP), enhanced green fluorescent protein (eGFP), and a Strep II tag (WSHPQFEK) were added to the C-terminus of the constructs. Site-directed mutagenesis was performed as previously described[83]. GluN3A chimeric constructs were generated by homologous recombination.

#### Mouse lines

Mice, GluN3A-KO and WT littermates, were housed and handled as previously described[31]. All experiments were performed in compliance with the French and European regulations (EU Directive 2010/63, French Law 2013-118, February 6th, 2013, authorization numbers #28867 and #44478).

#### eGlyR expression and purification

Recombinant baculovirus was produced using sf9 inset cells (ATCC CRL-1711) and followed the standard protocol of the Bac-to-Bac TOPO Expression System (Invitrogen A11339). Suspended HEK293S GnTI$^{-}$

cells (ATCC CRL-3022) at a concentration of 4.0 x $10^6$ cells/mL were infected using P2 virus. HEK293S GnTI⁻ cells were cultured in Freestyle 293 expression medium supplemented with 2% FBS and incubated at 37°C with shaking. 12 hours post-infection, 10 mM sodium butyrate was added into the culture medium to enhance protein expression. The cells were subsequently incubated at 30°C for another 48 hours and collected by centrifugation at 7,000g for 20 minutes. The cells were suspended and sonicated in TBS buffer (150 mM NaCl and 20 mM Tris, pH 8.0) containing a protease inhibitor cocktail (1 mM PMSF, 0.8 µM aprotinin, 2 mM pepstatin A, and 2 µg/mL leupeptin). The suspension was then solubilized by supplementing with 1% (w/v) L-MNG and 2 mM CHS for 2 hours at 4°C. After ultracentrifugation at 40,000 rpm for 1.5 hours, the supernatant was collected and incubated with Streptactin beads for 1.5 hours. The resin was collected and washed with 10 column volumes of TBS buffer supplemented with 0.1% (w/v) LMNG, 2 mM CHS and 200 µM EDTA. The protein was eluted with TBS buffer supplemented with 0.1% (w/v) digitonin, 5 µM CHS, 200 µM EDTA and 5 mM d-desthiobiotin. To remove the fused GFP, GluN3A-containing receptors protein was digested with 3C protease (1:20 molar ratio) for 4h at 4°C before loaded onto a Superose increase 6 10/300 GL column. Finally, the protein was concentrated and loaded onto a Superose increase 6 10/300 GL column (GE Healthcare) equilibrated with TBS buffer supplemented with 0.1% (w/v) digitonin, 5 µM CHS and 200 µM EDTA. The peak fraction was collected and concentrated to ~4.5 mg/mL using a 100 kDa cut-off Amicon Ultra Centrifugal Filter (Millipore). To enhance disulfide cross-linking, the GluN1/GluN3A-P497C receptors were treated with 500 mM $Cu^{2+}$(1,10-phenanthroline)$_3$ for 2 hours prior to being loaded onto a Superose Increase 6 10/300 GL column. To increase protein yield, the GluN1 antagonist CNQX was added during expression of the GluN1/GluN3A-S892L-K895V receptor, followed by the addition of 200 nM CGP to occupy the same site during subsequent purification.

**Cryo-EM sample preparation**

Samples (2.5 - 3.0 µL) were supplemented with individual ligands prior to plunge freezing. For the Gly/Gly bound state in both the GluN1/GluN3A-LF$^{EM}$ receptor and the GluN1/GluN3A$^{EM}$ receptor, 1 mM glycine was supplemented. For the CGP-78608/Gly bound state in the GluN1/GluN3A-LF, GluN1/GluN3A-LV, GluN1/GluN3A-P497C, and the GluN1/GluN3A-(2B NTD) receptors, 500 µM CGP-78608 was pre-incubated for 1 hour before supplementing with 1 mM glycine. For the Apo/Gly bound state in GluN1/GluN3A-LF$^{EM}$ receptor, no additional glycine was supplemented. The samples were applied to glow-discharged Quantifoil 1.2/1.3 Au 300 mesh grids, blotted, and plunge-frozen in liquid ethane using an FEI Vitrobot at 8°C and 100% humidity. The grids were initially screened using an FEI Talos 120C microscope, with particles evenly distributed and clearly visible.

**Cryo-EM data collection and processing**

Cryo-EM data were collected on a 300-kV Titan Krios G4 electron microscope (Thermo Fisher Scientific) using Falcon4 cameras with a slit width of 10 eV energy filter. Image acquisition was performed using EPU software (Thermo Fisher Scientific) with a defocus range of -1.0 to -2.5 µm, and the data were dose-fractionated over 40-50 frames with a total dose of 60 electrons. Cryo-EM image stacks were aligned using MotionCor2 1.4.0[84]. Contrast transfer function (CTF) parameters were estimated by Gctf_v1.06[85]. Approximately 3000 particles were selected for reference-free 2D classification, and the resulting 2D class averages were used as templates for autopicking in CryoSPARC 4.0[86]. Multiple rounds of 2D classification were performed to discard poor-quality particles. After several rounds of 3D classification, two distinct NTD conformations were distinguished. With the LBD still exhibiting consistent conformations in both NTD states, we combined the particles and first performed non-uniform refinement, followed by local refinement using a mask of the LBD and TMD. All maps were postprocessed using a solvent mask, with B-factors estimated automatically. Resolutions were determined using the gold-standard Fourier shell correlation at a 0.143 threshold. Local resolution estimation was performed in CryoSPARC 4.0.

**Cryo-EM model building**

The initial model of rat GluN1/GluN3A was generated using SWISS-MODEL[87]. Ligand coordinates and geometry restraints were obtained with phenix.elbow in Phenix 1.20-4459. [88] Models were docked into the EM density map using Chimera X 1.6.1[89]. This model was then refined through manual adjustments in Coot, followed by real-space refinement in Phenix. The final refinement statistics were validated using the "comprehensive validation (cryo-EM)" module in Phenix. For the low resolution of eGlyR NTD, we performed rigid body fitting using the structure predicted by[90], followed by refinement in Phenix and manual adjustments in Chimera X 1.6.1. A summary of the model refinement statistics is provided in Tables 1, 4 and 5.

**Structural analysis**

For structural analysis, the NTD and LBD were separated into R1 and R2, and D1 and D2 lobes. The center-of-mass (COM) of each domain or lobe was generated by PyMOL 2.5.2 using the 'Center-of-mass' script, and the distance between the COMs was then measured. After superimposing the two heterodimers, the domain rotation angle was determined using the RotationAxis scripts in PyMOL. The relative distance of the α-Helix E in GluN2/3-containing subunits, which serves as a readout of the tension exerted by the LBD to open the channel gate, was measured by the distance between the COMs of GluN2A (residues 668-671), GluN2B (residues 669-672), and GluN3A (residues 662-786). The degree of LBD opening–closure is assessed by measuring dihedral angles connecting the Cα of I403, S688, V735 and A715 in GluN1 and of P124, E525, S309 and E169 in GluN3A, respectively. Extended Data Fig. 7: the center of the R1 lobe in GluN1, GluN3A, GluN2A, and GluN2B used the CA position of the central amino acid within the following sequences YAILVSH, SALLAFP, HGLVFGD, and QGVVFAD, respectively. Similarly, the center of the R2 lobe in GluN1, GluN3A, GluN2A, and GluN2B used the CA position of the central amino acid within YVWLVGE, LHWVLGD, FFWIVPS, and YTWIVPS, respectively. Structural illustrations generated either with Chimera X or Pymol (PyMOL Molecular Graphics System, Version 3.0 Schrödinger, LLC.)

**3D modeling and simulation**

Model building of full size eGlyR (including all unresolved loops and linkers but not the C-terminal Domain). We selected one model given by Alphafold-3 with good statistics, and similarity with our experimental CryoEM densities[91] (Please note that the pIDDT of the LBD-NTD linker residues were always <<70 in coherence with the high predicted flexibility of the region). Topology and linkers shape of the model were adjusted using ModLoop (https://modbase.compbio.ucsf.edu/modloop/). The model was converted in full atom elastic network and fitted on the corresponding active state densities of GoF1 eGlyR in CGP+Gly ssing iModFit.[61] iModfit parameters were chosen as: cutoff: 0.1; -n modes/steps range: 0.05; -i iteration 1000000. Modelling of eGlyR conformational transition were performed with two steps iModFit simulation (See also [8,61,62] for iModfit parameters tests on NMDAR structures). iModfit parameters of first step were chosen as: cutoff: 0.1; -n modes/steps range: 0.05; -i iteration 1000000; -e 0.05 to speed up convergence and the -nowrmsd option. Parameters for the second step were: cutoff: 0.1; -n 0.3; -1 1000000 and other options by default. Extraction of collective variables was made using VMD[92].

**Two-electrode voltage-clamp (TEVC) recordings on Xenopus oocytes**

*Xenopus laevis* eggs were used for heterologous expression of recombinant NMDA receptors. Oocytes were harvested and maintained as previously described[83]. Expression of recombinant NMDA receptors was obtained by oocyte nuclear co-injection of 46 nL of a mixture of cDNA (50 ng/µl at 1:1 ratio). TEVC recordings were performed 48-72 hours after injection. Currents were recorded using an oocyte Clamp Amplifier OC-725 (Warner Instruments) and pClamp 10.5 (Axon, Molecular devices). During the recording, cells were perfused with an external Ringer recording solution (in mM: 100 NaCl, 0.3 $BaCl_2$, 5 HEPES, 2.5 KOH, pH adjusted to 7.3 with NaOH) and clamped at -60 mV at room temperature.

**Whole-cell patch-clamp recordings in HEK cells**

Patch-clamp recordings were performed on HEK293T cells. HEK293T cells were cultured in Dulbecco's modified Eagle's medium (DMEM) supplemented with 10% fetal bovine serum (FBS) at 37°C with 5% $CO_2$. GluN1 and GluN3A or GluN1 and GluN2B plasmids were transfected into HEK293T cells using Lipofectamine 3000 at a 1:1ratio. Transfected cells were digested with trypsin and transferred onto coverslips 30 minutes prior to recording. The bath solution contained 140 mM NaCl, 2.8 mM KCl, 1 mM $CaCl_2$, 10 mM HEPES, 10 mM EGTA, and pH was adjusted to 7.3 with NaOH. The pipette solution contained 115 mM CsF, 10 mM CsCl, 10 mM HEPES and 10 mM EGTA. Drugs and agonists were applied using a multi-barrel solution exchanger (RSC 200; Bio-logic). Currents were sampled at 10 kHz and low-pass filtered at 2.9 kHz using a HEKA EPC10 amplifier and Clampex 10.6. Recordings were performed at -60 mV and at room temperature. Recordings with series resistance greater than 25 MΩ or changes exceeding 20% were discarded to ensure data reliability. The electrode resistance in the bath solution was maintained between 5 and 8 MΩ to minimize signal distortion.

**Whole-cell patch-clamp recording in acute brain slices**

*Ex vivo* electrophysiology on acute brain slices was performed as previously described[31]. In brief, juvenile (P8-10) or adult (~2 months old) mice were deeply anesthetized with isoflurane and decapitated. Brains were rapidly removed and immersed in ice-cold sucrose-based artificial cerebrospinal fluid (ACSF) containing (in mM): 86 NaCl, 2.5 KCl, 0.5 $CaCl_2$, 7 $MgCl_2$, 1.2 $NaH_2PO_4$, 25 $NaHCO_3$, 25 glucose and 75 sucrose, continuously bubbled with 95% $O_2$/5% $CO_2$. Acute 300-μm horizontal slices containing the ventral hippocampus were prepared using a 7000 SMZ-2 Vibratome (Campden Instruments Ltd, UK). Slices were incubated for 30 min at 34 °C in standard ACSF containing (in mM): 125 NaCl, 2.5 KCl, 2 $CaCl_2$, 1 $MgCl_2$, 1.25 $NaH_2PO_4$, 25 $NaHCO_3$ and 25 glucose, then maintained at room temperature. For recordings, slices were transferred to a chamber and continuously perfused with oxygenated ACSF (3–4 mL $min^{-1}$, 30–34 °C). Borosilicate glass pipettes (3–5 MΩ) were filled with an intracellular solution containing (in mM): 130 K-gluconate, 0.6 EGTA, 2 $MgCl_2$, 0.2 $CaCl_2$, 10 HEPES, 2 Mg-ATP and 0.3 $Na_3$-GTP (pH 7.3, 295–300 mOsm) for glycine puffs experiments, and (in mM): 120 $CsMeSO_3$, 10 HEPES, 4.6 $MgCl_2$, 10 $K_2$-creatine phosphate, 15 BAPTA, 4 $Na_2$-ATP, 0.4 $Na_2$-GTP, 0.05 4-AP and 10 TEA-Cl, for NMDA puffs experiments. Signals were recorded using a Multiclamp 700B amplifier, a Digidata 1440A acquisition board and pClamp 10.3 software (Molecular Devices, USA), filtered at 2 kHz and digitized at 10 kHz; series resistance was compensated up to 65%. Glycine (1 mM) or NMDA (1 mM) was locally applied via pressure ejection (1 s). For glycine puff experiments, bath and puff solutions contained bicuculline (10 μM), strychnine (20 μM), NBQX (10 μM), D-AP5 (50 μM) and TTX (200 nM) to isolate GluN1/GluN3A-mediated currents. For NMDA puff experiments, strychnine and D-AP5 were omitted. I–V relationships were normalized to the charge measured at +40 mV.

**Protocols and Data Analysis**

For measurements in the presence of CGP-78608, the compound was pre-applied before exposure to the agonist: at 500 nM in the bath for recordings in HEK cells, and preincubated at 200 nM for recordings in Xenopus oocytes[25]. CGP-78608 potentiation values were obtained by calculating the ratio of peak current size for individual cells $I_{CGP+Gly}$ / $I_{Gly}$, with $I_{Gly}$ the peak current in 100 μM glycine before application of CGP-78608 and $I_{CGP+Gly}$ the peak current in 100 μM glycine during the application of CGP.

MTSEA potentiation was calculated by computing the ratio $I_m$ / $I_a$, where $I_a$ is the peak current obtained with the agonist after 200 nM CGP-78608 pre-incubation and and $I_m$ is the plateau current reached after addition of 300 μM MTSEA on top of the 200 nM CGP-78608 + 100 μM glycine. Relative channel maximal

Po values for the different receptor constructs were obtained by calculating the ratio 1 / MTSEA-potentiation.
Agonist dose-response curves (DRC) with standard sigmoid shape were fitted with the following equation: $I_{rel} = 1 / (1 + (EC_{50} / [Ago])^{nH})$, where $I_{rel}$ is the relative mean current (measured current / maximal current), [Ago] the agonist concentration (here glycine), and $n_H$ the Hill coefficient. $EC_{50}$ and nH were set as free parameters.
For agonist bell-shaped DRC, the fitting was performed with the following equation: $I_{rel} = max / (1 + (EC_{50} / [Gly])^{nH1}) * (1-min / (1 + (IC_{50} / [Gly]))^{nH2}))$, where max is the maximal relative value of the activation component, and max * (1-min) the maximal relative value of the inhibitory component. In order to help fitting this 6-parameters equation, we used as seeding values the activation parameters (max, $EC_{50}$ and $n_{H1}$) obtained from a first round of approximate fit with a standard single-component Hill equation.
For high affinity mutants, we subtracted the contribution of tonic activation to the recorded current at each glycine concentration (see Extended Data Fig. 3b). Tonic activation by contaminant glycine was revealed and measured for each oocyte by application of 100 µM of pore blocker pentamidine on Ringer alone.
For QRN site analysis ($MgCl_2$ DRC and I/V curves), we used GluN1/GluN3A receptors carrying the GluN1-F4984A mutation in order to prevent receptor desensitization (Awobuluyi, Grand et al). $MgCl_2$ inhibitory DRC were performed in saturating agonist (100 µM glycine) and analyzed with the following equation: $I_{rel} = Max + ((1-Max) / (1 + (IC_{50} / [MgCl_2])^{nH})$, where Max is the maximum inhibition at saturating $MgCl_2$ concentration, $I_{rel}$ is the relative mean current (measured current / maximal current), and $n_H$ the Hill coefficient. $IC_{50}$ and nH were set as free parameters. I/V curves of NMDARs were performed between -100 and +50 mV in saturating agonist (100 µM glycine). All I/V curves were normalized to the current value recorded at +25 mV. Data analysis was performed using Clampfit 10.6 (Axon Instruments) and Igor Pro 6.11 (WaveMetrics).

**Pharmacological Reagents**
CGP-78608 (Tocris) was prepared as a stock solution at 25 mM in 2.2 eq. NaOH solution. In electrophysiological experiments, CGP-78608 was applied either at 50 nM (slightly above $EC_{50}$) or at 200 nM (saturation) before agonist perfusion[25]. Aminoethyl-methanethiosulfonate (MTSEA; Toronto Research Chemicals Inc.) was prepared as a stock solution at 100 mM in DMSO. Pentamidine isethionate salt (Pentamidine, Sigma) was prepared at a stock concentration at 10 mM in bi-distilled water.

**Statistical Analysis**
Unless otherwise indicated, data are presented as mean ± standard deviation of the mean (SEM). Non-parametric tests were used to assess statistical significance. When comparing two conditions, the non-parametric Mann Whitney test was employed, while for more than two groups the non-parametric Kruskal-Wallis test was employed, with Dunn's multiple comparisons post hoc test. Statistical significances are indicated with *, **, *** when p- values are below 0.05, 0.01 and 0.001 respectively. n.s. indicates non-significant.

#### Extended Figure legends

**Extended Data Fig. 1: eGlyR constructs used for single-particle cryo-EM imaging exhibit WT-like properties**

**(a-c)** Representative current traces of NMDARs upon application of agonist alone (left) and with 500 nM CGP-78608 (right). The constructs expressed and recorded in HEK293 cells are the following: (A) wild-type (WT) GluN1/GluN3A receptors with glycine as agonist, (B) GluN1/GluN3A-(2B NTD) chimeric receptors (same conditions as in A) and (C) GluN1/GluN2A, with glutamate and glycine as agonists.

**(d)** Glycine sensitivity of wild-type GluN1/GluN3A and chimeric GluN1/GluN3A-(2B NTD) receptors in the presence of CGP-78608 (500 nM). The $EC_{50}$ values are 21.5 ± 1.5 µM [n = 4] for GluN1/GluN3A and 26.3 ± 5 µM [n = 4] for GluN1/GluN3A-(2B NTD) receptors.

**Extended Data Fig. 2: Particle processing and cryo-EM data for GluN1/GluN3A$^{EM}$ and GluN1/GluN3A-(2B NTD)$^{EM}$ receptors**

**(a)** Left: Representative current traces of wild-type GluN1/GluN3A and GluN1/GluN3A$^{EM}$ receptors activated by application of glycine alone (100 µM) and with CGP-78608 (500 nM). The responses are normalized to the peak current amplitude. The recordings were performed in HEK293 cells. Right: Time course ($\tau_{Inact}$) of current inactivation obtained by single-exponential fit. Error bars indicate mean ± SEM; n = 3 - 5.; **p < 0.01, ***p < 0.001, Student's t-test.

**(b,d)** Fluorescence size-exclusion chromatography of the purified GluN1/GluN3A$^{EM}$ (B) and GluN1/GluN3A-(2B NTD)$^{EM}$ (D) receptors. The inserted panel in (B) shows Coomassie Blue staining of the purified GluN1/GluN3A$^{EM}$ protein, with two bands corresponding to GluN1$^{\Delta CTD}$-GFP at 124.9 kDa and GluN3A$^{\Delta CTD}$-GFP at 137.2 kDa. The inserted panel in (D) shows Coomassie Blue staining of the purified GluN1/GluN3A-(2B NTD)$^{EM}$ receptor, with two bands indicating GluN1$^{\Delta CTD}$ at 96.2 kDa and GluN3A-(2B NTD)$^{\Delta CTD}$ at 97.6 kDa.

**(c,e)** Pipelines for single-particle analysis and reconstruction of GluN1/GluN3A$^{EM}$ receptors in glycine bound state (see Fig. 1d) and GluN1/GluN3A-(2B NTD)$^{EM}$ receptor in CGP-78608/glycine bound state (see Fig. 2e). Representative micrograph is illustrated with a scale bar of 300 Å. Particles with clear characteristic orientations from 2D class average are selected. The classes with similar conformation (in dash line square) are merged for the UN-refinement with C2 symmetry. Cryo-EM density map are colored based on the local resolution estimation by CryoSPARC. Fourier shell correlation curves (FSC) for cryo-EM density maps are plotted with (in blue) or without (in purple) masking.

**Extended Data Fig. 3: Gain-of-function phenotypes (GoF1) at the LBD GluN1-GluN3A dimer interface**

**(a)** Glycine dose-response curves (DRC) from peak currents of WT GluN1/GluN3A and GluN1/GluN3A-LF receptors in presence or absence of CGP-78608 (200 nM). See Table 2 for values of $EC_{50}$, Hill slope ($n_H$) and statistics.

**(b)** Quantification of tonic currents in GluN1/GluN3A-LF receptors. Right: representative current trace of GluN1/GluN3A-LF mutant receptors revealing the presence of tonic current (i.e. in nominal 0 glycine) inhibited by pentamidine (100 µM), a channel blocker of eGlyRs. Quantification of tonic currents (pentamidine effect) in relation to currents obtained in saturating glycine alone (100 µM, white bar) or in glycine plus CGP-78608 (200 nM; striped bar).

**(c)** GoF1 drastically reduces CGP-78608 potentiation. Quantification of the potentiation induced by CGP-78608 (200 nM) of WT and GoF1 receptors activated by 100 µM glycine. Inset: representative current trace of GoF1 in glycine alone and with CGP-78608. Note that glycine alone induces a reduction of the steady-state current above the baseline.

**(d)** Dose-response curve of CGP-78608 potentiation of GluN1/GluN3A-LF receptors in the presence of 100 µM glycine. Currents obtained in Xenopus oocytes were measured at the peak. CGP-78608 $EC_{50}$ = 33.1 ± 1.2 nM, $n_H$ = 1.41 (n = 6).
**(e)** Increasing hydrophobic contribution[93] of substituted resides correlates with increased glycine sensitivity (similarly than for the Po, see Fig. 2d). Each data point relates to specific receptor mutants corresponding at the GluN3A-S892-K895 position: GluN3A-S892L-K895F (LF), GluN3A-S892L-K895V (LV), GluN3A-S892L-K895I(LI) and GluN3A-S892L-K895W(LW). The lines represent a linear regression fit.
**(f-g)** Representative current traces of currents for wild-type GluN1/GluN3A and GluN3A-S892L-K895F(LF) receptors activated by glycine alone (A) and with GCP-78608 (500 nM, B). The recordings were performed in HEK293 cells. The corresponding dose-response curves are shown in the right panels.

**Extended Data Fig. 4: Particle processing and cryo-EM data for GluN1/GluN3A-$LV^{EM}$ receptors**
**(a)** Left panel: Size-exclusion chromatography (SEC) of purified GluN1/GluN3A-$LV^{EM}$ receptors. Right panel: Coomassie Blue staining of the purified protein, showing two bands corresponding to GluN1$^{\Delta CTD}$-GFP (124.9 kDa) and GluN3A$^{\Delta CTD,S892L,K895V}$-GFP (137.2 kDa).
**(b)** Pipelines of single-particle analysis and reconstruction of GluN1/GluN3A-$LV^{EM}$ receptor structure in Glycine and CGP-78608 (See Fig. 2f). Representative micrograph is illustrated with a scale bar of 300 Å. Particles with clear characteristic orientations from 2D class average are selected. Topaz was used to select 2D classes with disrupted LBDs, revealing that the NTDs were uncrossed and the LBDs displayed four-fold symmetry at low resolution. The classes with similar LBD-TMD conformation (in dash line square) were merged for the UN-refinement with C2 symmetry. Cryo-EM density map were colored based on the local resolution estimation by CryoSPARC. Fourier shell correlation curves (FSC) for cryo-EM density maps were plotted with (in blue) or without (in purple) masking. Local electron densities and fitted atomic models are shown for the TMD regions of the GluN1 and GluN3A subunits.

**Extended Data Fig. 5: Particle processing and cryo-EM data for GluN1/GluN3A-$LF^{EM}$ receptors**
**(a)** Left panel: Size-exclusion chromatography (SEC) of purified GluN1/GluN3A-$LF^{EM}$ receptors. Middle panel: Concentration of the peak fraction from purified GluN1/GluN3A-$LF^{EM}$ receptors, followed by fluorescence size-exclusion chromatography (FSEC) analysis. Right panel: Coomassie Blue staining of the purified protein, showing two bands corresponding to GluN1$^{\Delta CTD}$ (96.2 kDa) and GluN3A$^{\Delta CTD,S892L,K895F}$ (108.5 kDa).
**(b,e,h)** Pipelines of single-particle analysis and reconstruction of GluN1/GluN3A-$LF^{EM}$ receptor structure in three different conditions (See Fig. 2g). Representative micrograph is illustrated with a scale bar of 300 Å. Particles with clear characteristic orientations from 2D class average are selected. The classes with similar conformation (in dash line square) are merged for the UN-refinement with C2 symmetry. Classifications on LBDs and NTDs were performed by using soft masks on those regions or doing particle subtraction.
**(c,f,i)** Cryo-EM density map colored based on the local resolution estimation by CryoSPARC.
**(d,g,j)** Fourier shell correlation curves (FSC) for cryo-EM density maps with (in blue) or without (in purple) masking.

**Extended Data Fig. 6: Characterization of GoF2 mutant receptors**
**(a)** CGP-78608 induced potentiation of GluN3A NTD-LBD linker cysteine mutant receptors and their controls (A or S mutations). Results for wild-type (WT) GluN1/GluN3A receptors are also shown. The recordings were performed Xenopus oocytes. Cysteine mutants from GluN3A-P497C to GluN3A-Q501, each individually, exhibit clear GoF phenotypes but not mutants GluN3A-L513C and GluN3A-H514C (closer from the LBD – see alignment in Fig. 3a) which display WT-like phenotypes.

**(b)** GoF2 mutant receptors exhibit large and stable response with Glycine but only modest change in glycine sensitivity compared to GluN3A WT. Top: TEVC current traces of GluN1/GluN3A-Q501C upon application of increasing concentrations of glycine. Recorded in xenopus oocyte. Bottom: Glycine dose-response curves for WT and GluN1/GluN3A-Q501C receptor in presence or absence of CGP-78608 (200 nM). Refer to Table 2 for values of $EC_{50}$ and Hill slope (nH).
**(c)** Representative current traces of glycine sensitivity of GluN1/GluN3A-Q501C receptors recorded in HEK293 cells. Note the greatly reduced desensitization compared to WT receptors (see Fig. 2b and Extended Data Fig. 3f; see also panel f below).
**(d)** Reduced desensitization in the GoF2 GluN3A NTD-LBD linker cysteine mutant receptors compared to WT receptors. Receptors were activated by 100 µM glycine alone. Recorded in HEK293 cells.
**(e)** Formation of disulfide crosslinks in GoF2 mutant receptors. Effect of the reducing agent DTE on current amplitudes of GluN1/GluN3A-Q501C receptors and their controls. Note the opposite direction of the DTE effect between the two set of receptors.
**(f)** Western blot from Xenopus oocytes[62,94] expressing different eGlyR constructs and revealed with an anti-GluN3A antibody (catalog #07-356, Sigma-Aldrich). showing the presence of both GluN3A monomer and GluN3A dimer band in the five Go-F2 cysteine mutants but not in WT receptors and non-injected oocytes (NI). Dashed red square: zoom of the GluN3A dimer band region with enhanced contrast.

**Extended Data Fig. 7: High conformational dynamics of eGlyR NTDs**
**(a)** Side view (top) and top-down view (bottom) of the extracellular domain (ECD) densities of (left to right): GluN1/GluN3A-$LF^{EM}$ receptors in NTD crossed and uncrossed conformations, GluN1/GluN3A-(2B $NTD)^{EM}$ and GluN1/GluN2B receptors. GluN1, GluN3A, and GluN2B in grey, red, and blue, respectively. Note the major rearrangements of the NTDs between the two conformations.
**(b)** Normal-mode based trajectory between the crossed and non-crossed states. The crossed-state model of GluN1/GluN3A-$LF^{EM}$ ECD is shown in ribbon with GluN1 in light grey and GluN3A in dark grey. Red arrows indicate the direction of ECD motion during the trajectory.
**(c)** Class analysis of NTD conformations in GluN1/GluN3A-$LF^{EM}$ receptor densities (CGP/Gly conditions). Ten classes were defined, with four different NTD conformations: crossed (black, 6 classes), uncrossed (brown, 1 class), and two distinct intermediate conformations intermed 1 (purple, 2 classes) and intermed 2 (green, 1 class). The five collective variables (three distances and two angles) used to quantify the conformational differences are shown as single (1 class) or mean (several classes) values.
**(d)** Evolution of the RMSD calculated for each step of the simulated trajectory (see B) against the four experimentally determined NTD conformations (see C). Note that intermed 1 (purple) and intermed 2 (green) show a minimum at step 29 and 53 of the simulation, respectively.
**(e)** Evolution of the five collective variables (from bottom to top, see C) of GluN1/GluN3A-$LF^{EM}$ receptors when transiting from the crossed to uncrossed conformations. Experimentally-determined data (symbols, color code as in C) and simulated trajectory (lines) are superimposed. In red on the right: Values of the same collective variables for GluN1/$GluN3A^{EM}$ receptors in the desensitized-like state (from Extended Data Fig. 2c).

**Extended Data Fig. 8: Particle processing and cryo-EM data for GluN1/GluN3A-$PC^{EM}$ receptors**
**(a)** Left panel: Size-exclusion chromatography (SEC) of purified GluN1/GluN3A-$P497C^{EM}$ receptors. Middle panel: SEC profile of purified GluN1/GluN3A-$LV^{EM}$ receptors in the presence of the oxidant Cupper phenanthroline. Right panel: Coomassie Blue staining of the purified protein, showing two bands corresponding to $GluN1^{\Delta CTD}$-GFP (124.9 kDa) and $GluN3A^{\Delta CTD,P497C}$-GFP (137.2 kDa).
**(b)** Pipelines of single-particle analysis and reconstruction of GluN1/GluN3A-$PC^{EM}$ receptor structure in the presence of Glycine and CGP-78608 (See Fig. 3e). Representative micrograph is illustrated with a scale bar of 300 Å. Particles with clear characteristic orientations from 2D class average are selected. The

classes with similar conformation (in dash line square) are merged for the UN-refinement with C2 symmetry. Classifications on the LBD-TMD region were performed by using soft masks on those regions or doing particle subtraction. In the middle, Cryo-EM density map colored based on the local resolution estimation by CryoSPARC. On the right, Fourier shell correlation curves (FSC) for cryo-EM density maps with (in blue) or without (in purple) masking.

**Extended Data Fig. 9: eGlyR gating mechanism**
Measurements of LBD opening angles in all available structures of GluN3A containing receptors. Open triangles are for GluN1 subunits and filled triangles are for GluN3A subunits. For comparison purposes, angles extracted from previously published receptor structures are included: 'CC' for GluN1/GluN3A CNQX/Gly structure (8USW)[95] , 'Crystal Apo' for Apo GluN3A LBD (4KCD)[96] and Apo G&luN1 LBD (4KCC)[96], and 'Crystal Gly' for glycine-bound GluN3A LBD (2RC7)[18] and glycine-bound GluN1 LBD (4NF8)[97].

**Extended Data Fig. 10: Structural comparison of the LBD and NTD organization between GluN1/GluN3A eGlyRs and AMPA/kainate receptors**
**(a)** Top: Top-down views of the LBD tetramer arrangement in : pre-active GluN1/GluN3A-PC and desensitized GluN1/GluN3A-WT receptors (this study); active AMPA (PDB 8C1P)[51] and desensitized AMPA (PDB 8P3W)[50] and active kainate (PDB 9B36)[51] and desensitized kainate (PDB 9B38)[51] receptors. Bottom: Zoom in the respective LBD dimer conformation.
**(b)** Comparison of the structural arrangements between NTD and LBD layers in eGlyRs, AMPA and kainate receptors. Conformational differences between subunits are quantified by vector angles connecting the COMs of the R2 and R1 lobes of the NTDs, and the D1 and D2 lobes of the LBDs. The distances between domains are measured using their COMs (see Methods).

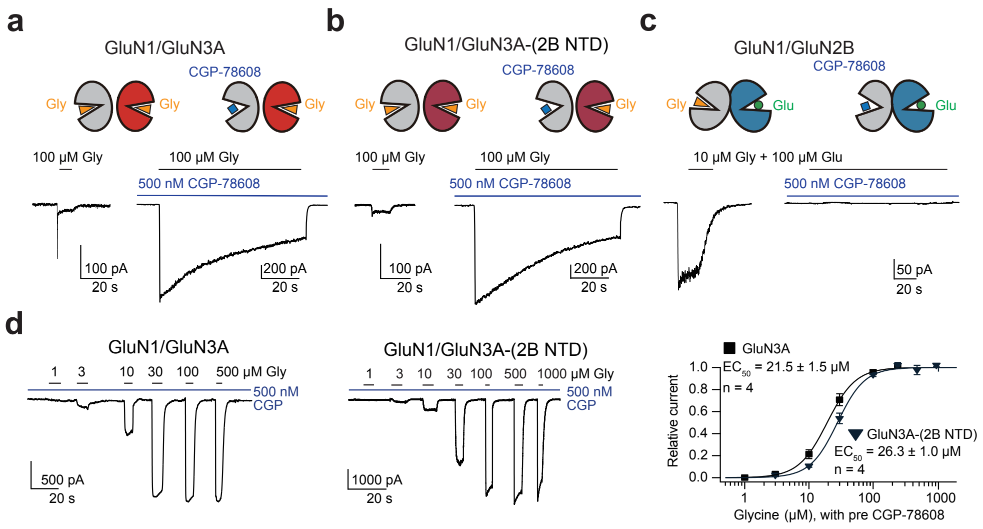


### Extended Data Fig.1

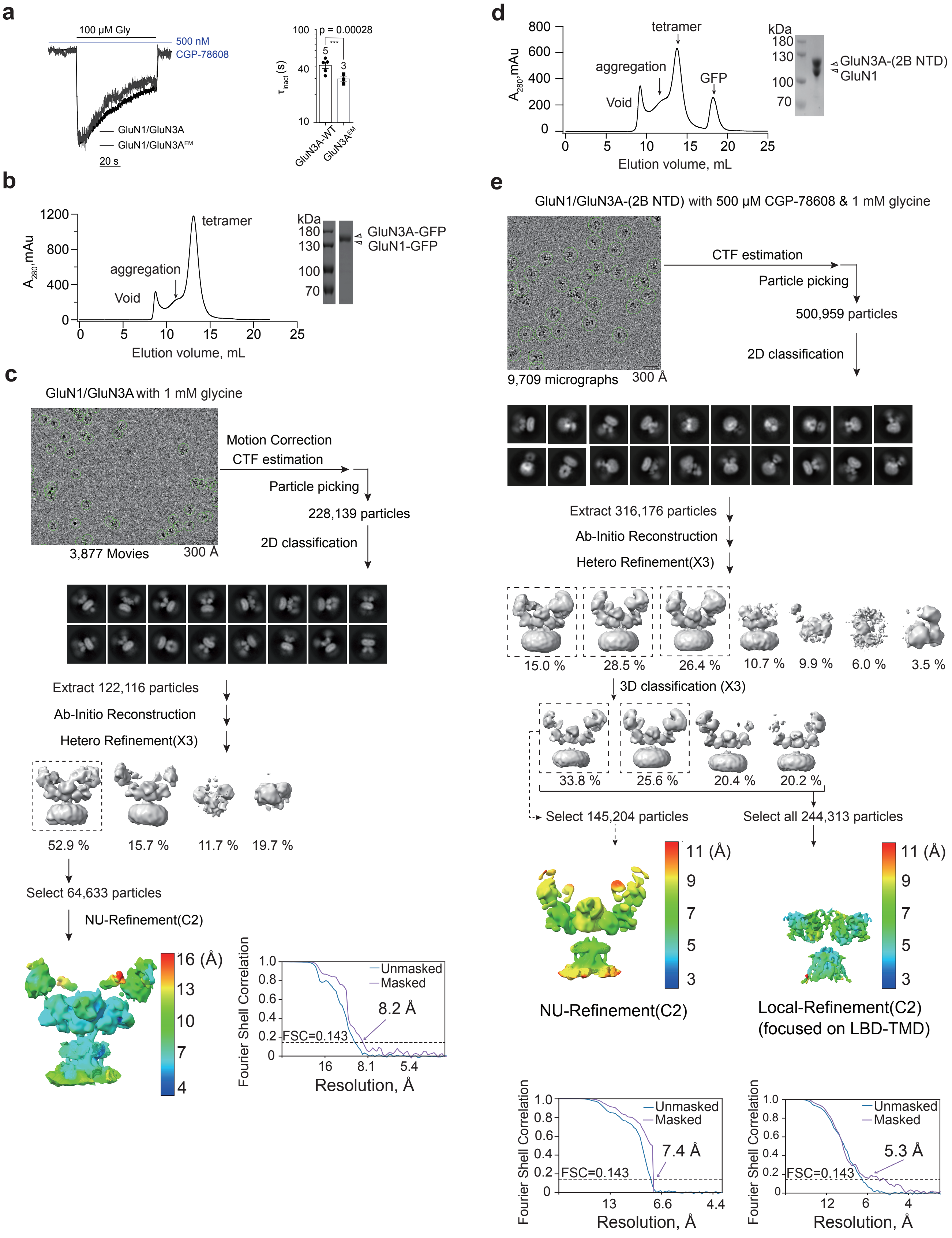


### Extended Data Fig.2

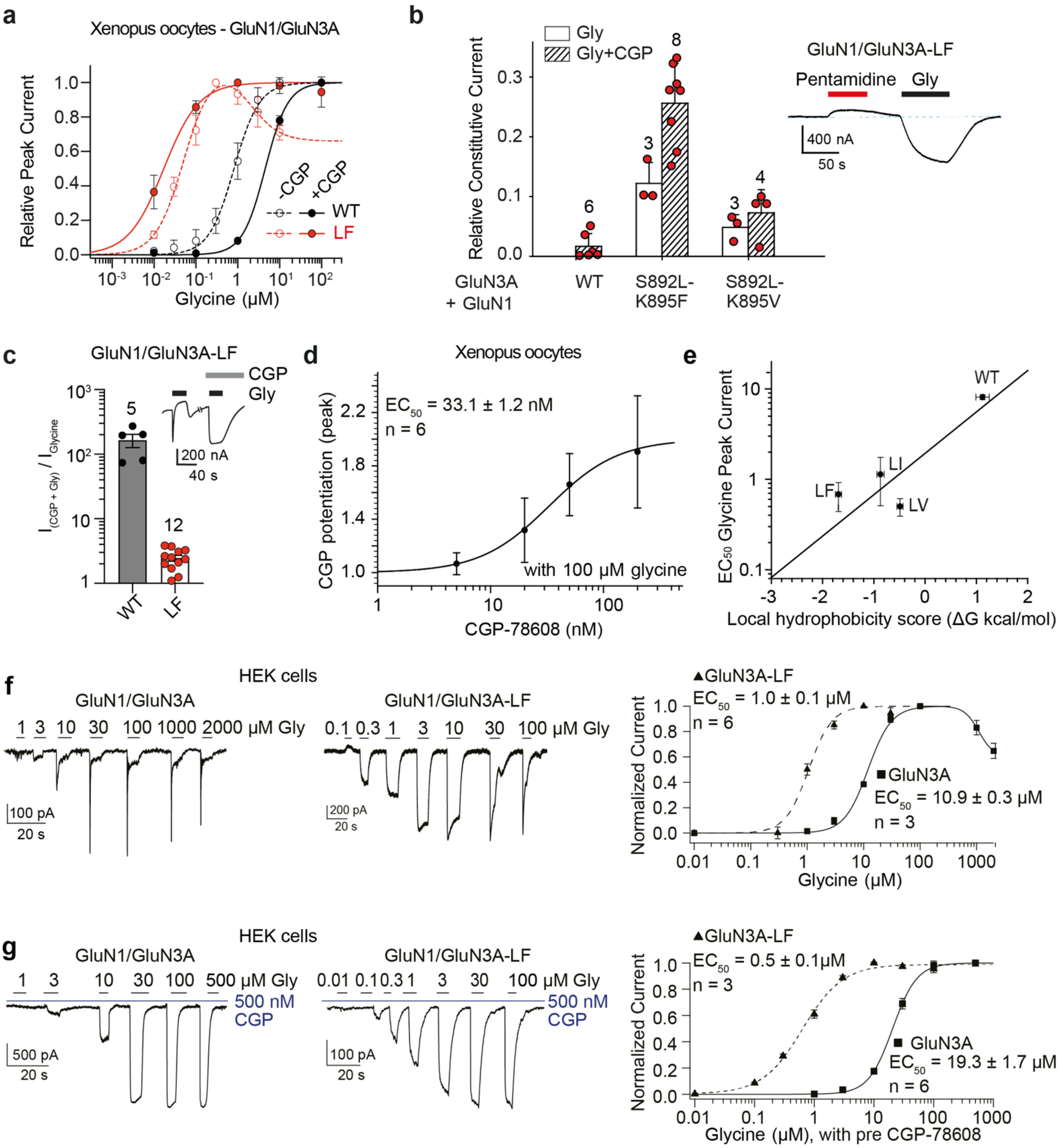


**Extended Data Fig.3**

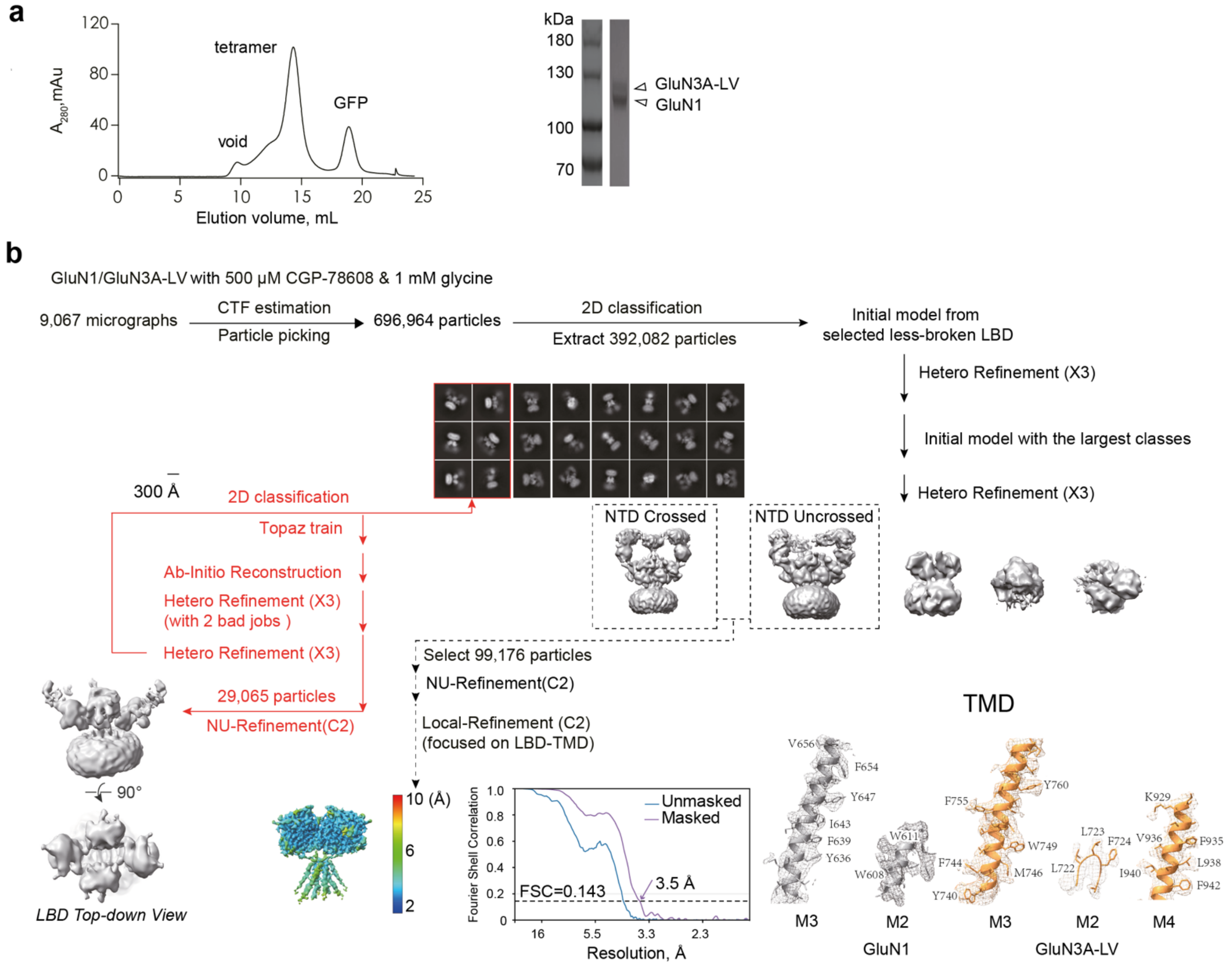


### Extended Data Fig.4

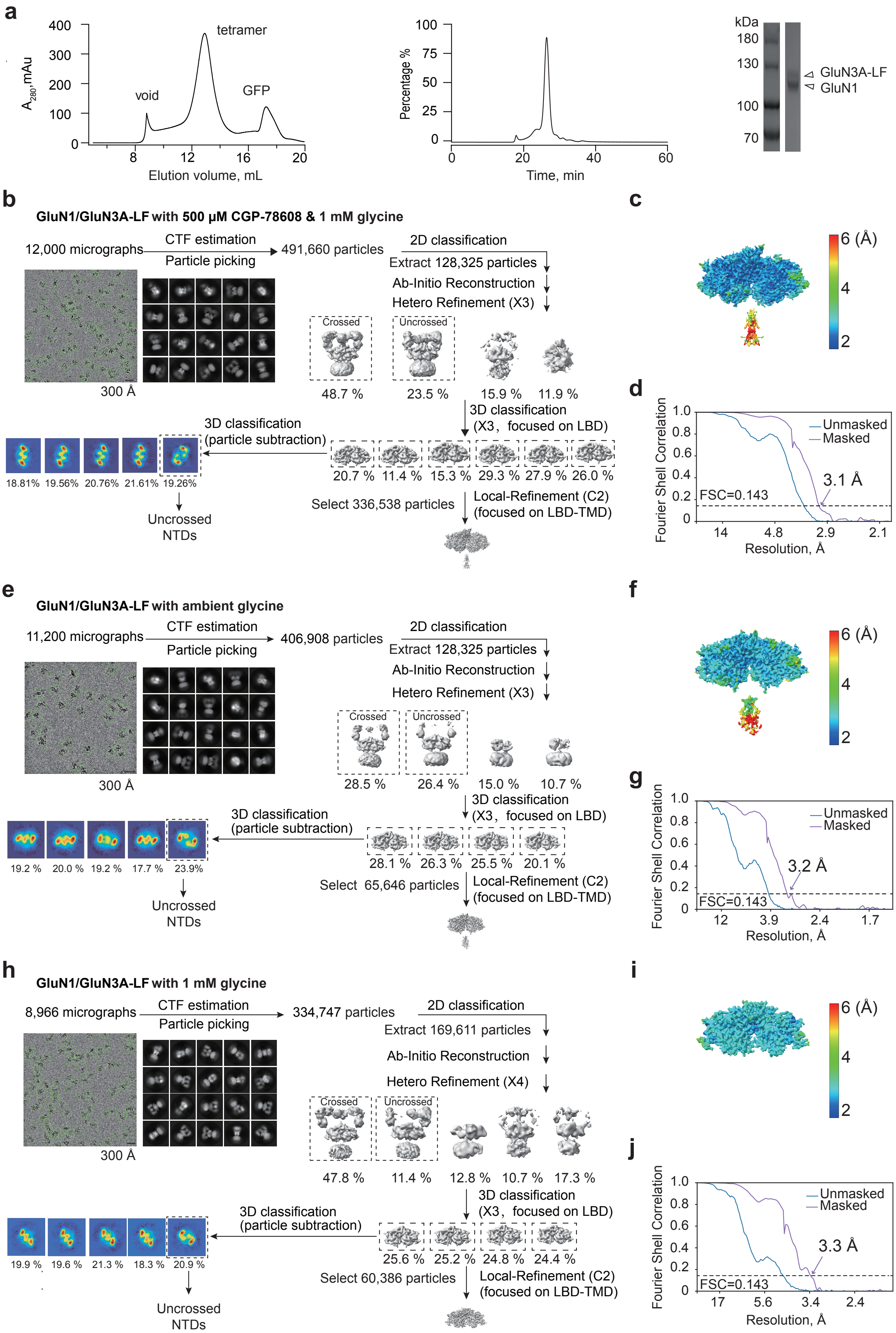


### Extended Data Fig.5

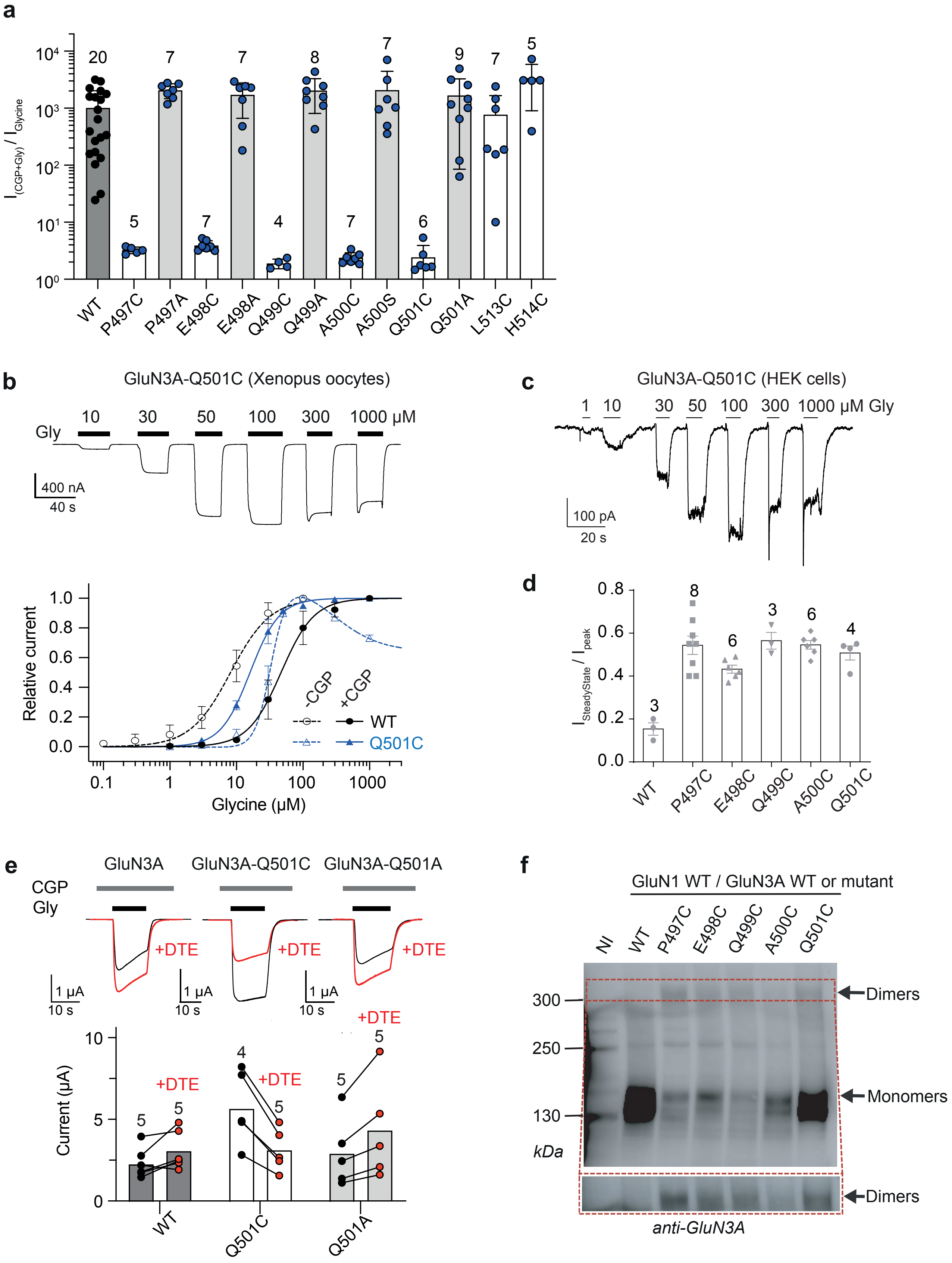


### Extended Data Fig.6

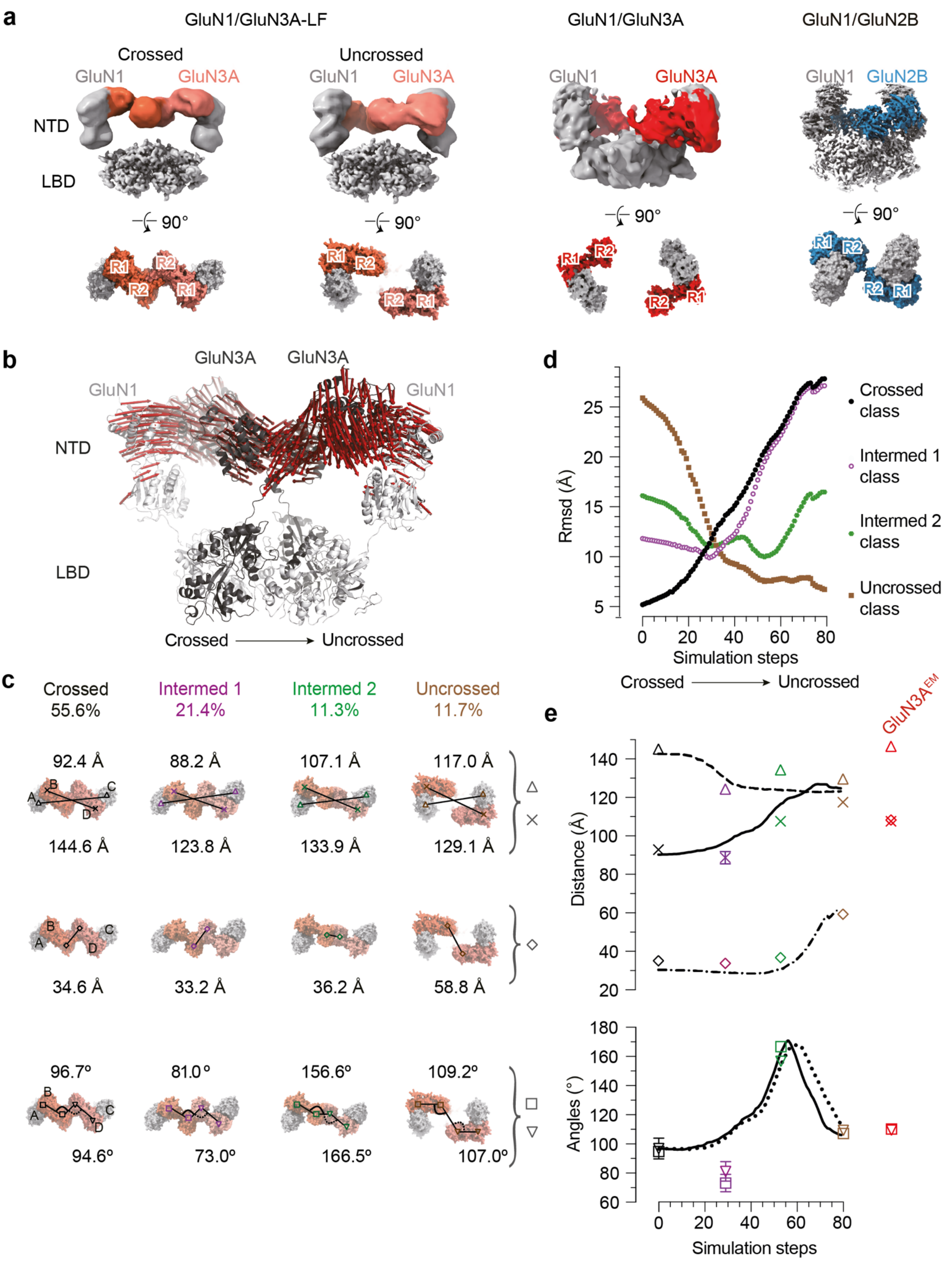


**Extended Data Fig.7**

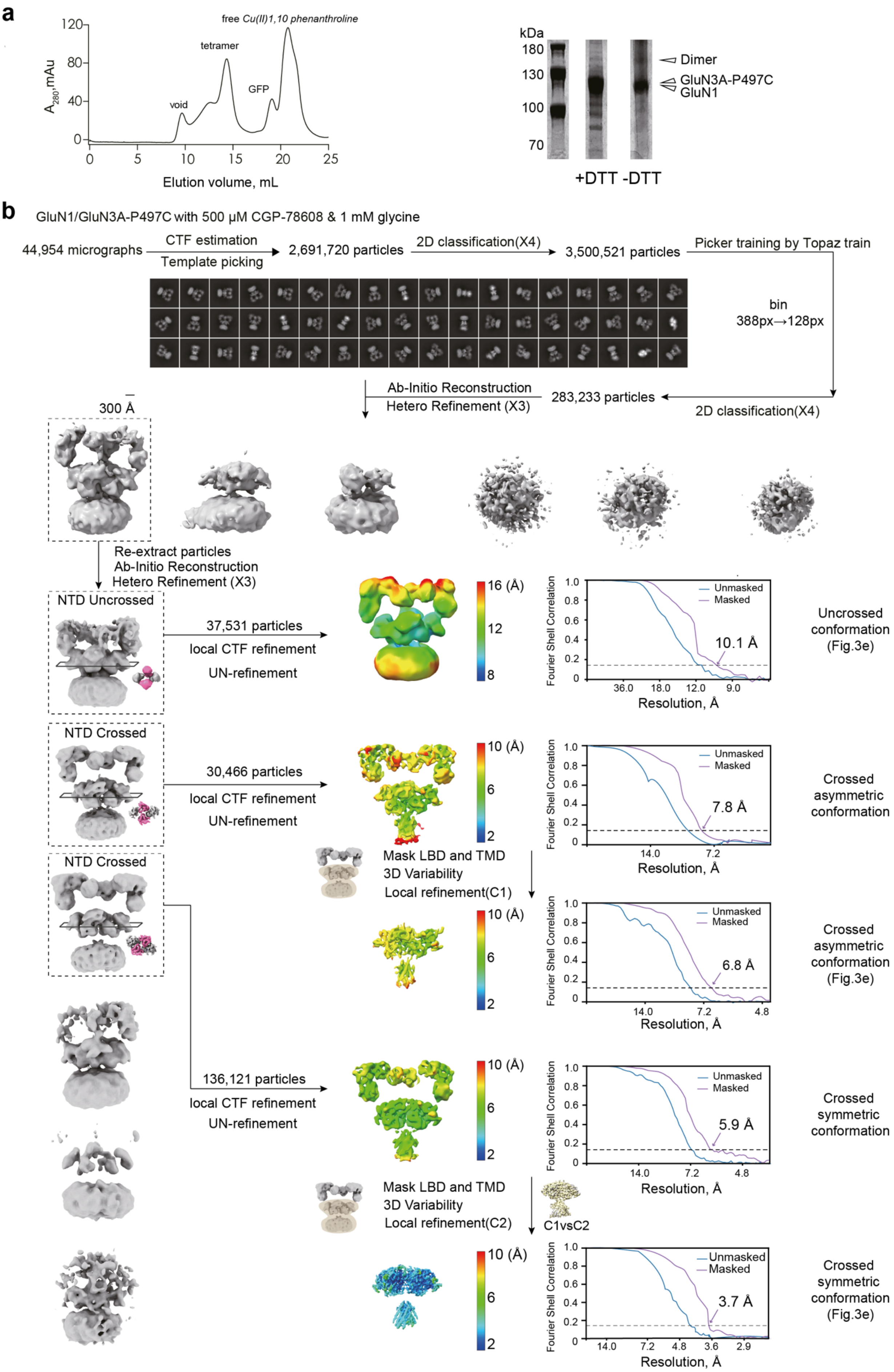


**Extended Data Fig.8**

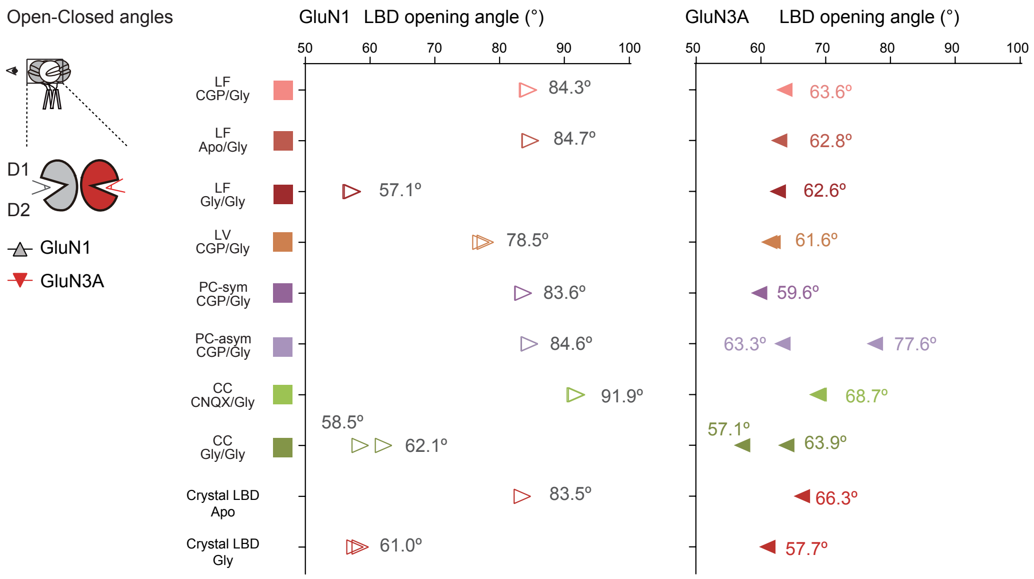


### Extended Data Fig.9

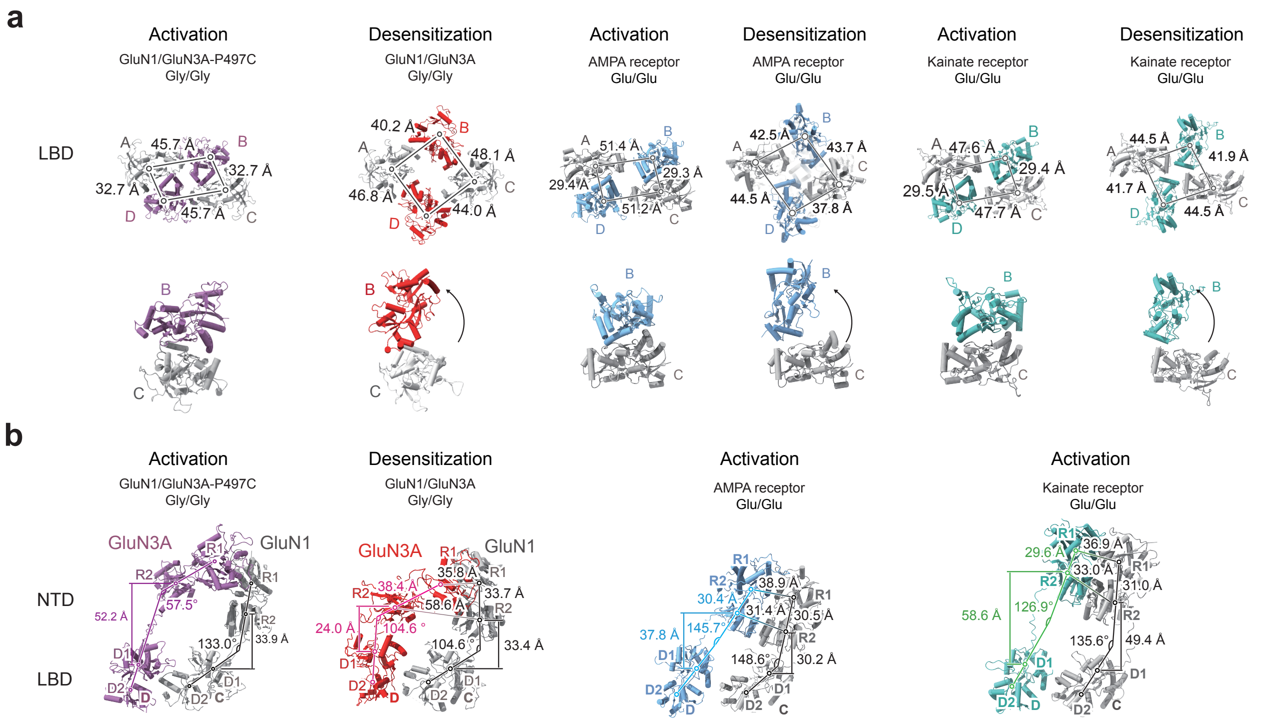


### Extended Data Fig.10

#### References (Methods and Extended Data)
